# An ARF2-GRF5 module regulates chloroplast biogenesis through GLK1 and GLK-independent mechanisms as part of a genetic network

**DOI:** 10.1101/2025.06.29.662209

**Authors:** Priyanka Mishra, Julian M Hibberd, Enrique López-Juez

**Affiliations:** Department of Biological Sciences, Royal Holloway University of London, Egham TW20 0EX, United Kingdom; Department of Plant Sciences, University of Cambridge, Downing Street, Cambridge CB2 3EA, United Kingdom

**Keywords:** Chloroplast, plastid, Arabidopsis, GLK, ARF2, GRF5, photosynthetic differentiation, suppressor mutation

## Abstract

- Our current understanding of the regulation of chloroplast development lags behind that of the process itself. The GOLDEN2-LIKE (GLK) factors drive the expression of photosynthesis-associated nuclear (PhAN) genes. Simultaneous loss of GLK1 and GLK2 in Arabidopsis plants leads to pale-green, incompletely-developed chloroplasts. This highlights the GLKs’ role but also the involvement of other components.
- We hypothesised that further regulators could be found by searching for suppressor mutations of a severe *glk1* knock-down (KD) *glk2* knock-out (KO) mutant. We identified *suppressor of glks* (*sgl*) *2*, *sgl2*.
- The *sgl2 glk1 glk2* triple mutant showed significant increases in chlorophyll, expression of PhAN genes, chloroplast thylakoid stacking, and cellular volume of chloroplasts. *sgl2* was caused by loss of function of *AUXIN RESPONSE FACTOR2*, *ARF2*. *arf2^sgl2^* boosted chloroplast content as a single mutant, and dramatically suppressed the chloroplast delay of the phytochrome-deficient *long hypocotyl 1* (*hy1*) mutant. Overexpression of *Growth Regulating Factor 5* (*GRF5*), which is negatively regulated by ARF2, also rescued chlorophyll levels in *glk1 glk2*, confirming a role for ARF2 and GRF5 as a transcription module. We observed in *glk1 glk2* low but detectable expression of *GLK1*, which was elevated in *arf2^sgl2^ glk1 glk2*, and saw reduced suppression by *arf2^sgl2^* of the phenotype of a separate *glk1*-2 *glk2*-2 mutant carrying two KO alleles, revealing partial epistasis. Presumed orthologs of *ARF2* and *GRF5* are expressed very early in developing cereal leaves, well ahead of the bulk of *GLK* expression during the greening stage.
- Our results place ARF2 as a negative switch modulating *GRF5* early and upstream, whether directly or indirectly, of GLK factors, but also acting independently from them. This reveals a hierarchy of transcription factor action and feed-forward regulation in chloroplast biogenesis during leaf development.

## Introduction

The most substantial source of energy and oxygen support of life on this planet comes from organisms carrying out oxygenic photosynthesis, a process that is nearly as ancient as the existence of life (Sakamoto et al., 2008; Björn and Govindjee, 2009; Moody et al., 2024). This life-supporting process is performed in eukaryotes by chloroplasts, cell organelles specialised in capturing sunlight. Chloroplasts, to a greater extent than mitochondria in mammals, vary in extent of development and final function across multiple cell types (Jarvis and López-Juez, 2013; Sierra et al., 2023). Although our knowledge of chloroplast development and function is substantial, that of the regulation of the biogenesis of chloroplasts is very limited. The "Green Revolution" involved expanding irrigation and fertilizer use, and developing high-yield crop varieties that benefit from the extra fertilizer. Most of those improvements have been maximized. However, enhancing photosynthesis remains a promising and underexplored avenue (Eckardt et al., 2023; Croce et al., 2024). One of the potential sustainable tools for enhancing photosynthesis is to increase or adjust chloroplast volume in leaf cells. Identifying regulators of chloroplast development is key to efforts to engineer the "chloroplast compartment size" (Li et al., 2020; Cackett et al., 2022).

Our current understanding of early chloroplast development and its regulation, particularly during the phase when plastids are filling the cellular space, is very limited (Loudya et al., 2021). Regulation of early plastid development occurs at a time of intense cellular proliferation, when plant stem cells enter, in leaf primordia, a phase akin to a “transient amplifying” stage in organ development, and this involves a multitude of auxin and cytokinin-dependent responses (Braybrook and Kuhlemeier, 2010; Tsukaya, 2013; Mohammed et al., 2018). Genes associated with hormone signalling act even before the expression of the bulk of transcription factors (TFs) essential for chloroplast development (Loudya et al., 2021; Cackett et al., 2022). Light perception, a main cue, promotes the function of transcription factors for photomorphogenesis, including chloroplast development (Quevedo et al., 2025). For example, LONG HYPOCOTYL 5 (HY5) regulates chloroplast development in the light (Martín et al., 2016) by promoting the expression of key regulators of chloroplast biogenesis such as the GOLDEN2-LIKE (GLK1 and GLK2) and the GATA family - including GNC (GATA nitrate-inducible carbon metabolism-involved) and CGA1 (cytokinin-responsive GATA factor 1) (Chiang et al., 2012; Zubo et al., 2018; Zhang et al., 2024). Conversely, phytochrome interacting factors (PIFs) repress plastid development in the dark, working antagonistically to HY5 (Richter et al., 2010; Cackett et al., 2022). In addition, GRF5 (GROWTH REGULATING FACTOR5) is reported to impact chloroplast division, enhances photosynthesis, and extends leaf lifespan, functioning partially redundantly with eight other members of the GRF family in Arabidopsis (Kim et al., 2003; Horiguchi et al., 2005; Vercruyssen et al., 2015). Two newly identified RR-type myoblastoma-related (RR-MYB) TFs also promote chloroplast biogenesis in *Marchantia* and Arabidopsis under light conditions (Frangedakis et al., 2024). Loss-of-function of these light-induced TFs results in reduced chlorophyll biosynthesis and produces pale plants, highlighting a positive role in chloroplast development. Adapting this understanding to generate tools for enhancing photosynthesis will involve generating overexpressor lines. Identification of negative regulators of chloroplast development for the above-mentioned factors would enhance our understanding of their specific role and interaction in overall chloroplast-making.

GLK1 and GLK2 are crucial for pigment biosynthesis and chloroplast development (Fitter et al., 2002; Waters et al., 2008; Waters et al., 2009; Wang et al., 2013). Their disruption only partially affects chloroplast biogenesis, suggesting a parallel role for other known (RR-MYBs) and unknown genes. Identifying specific early chloroplast regulators is challenging due to the multitude of genetic circuitry involved in early cellular development across different cell types. In a highly-resolved wheat leaf model of proplastid to chloroplast differentiation, GLK1 homologs are primarily expressed during the later, greening phase (Loudya et al., 2021). This implies a limited role in earlier plastid development and the need for discovery of early-acting regulators (Wang et al., 2017; Loudya et al., 2021). We sought to identify additional regulators of chloroplast development (López-Juez and Hills, 2011) in the absence of GLK master regulators. The pale phenotype of the *Arabidopsis glk1 glk2* double mutant constitutes an advantageous system for identifying independent greening-inducing chloroplast regulators using genetic tools.

We screened an ethyl methanesulfonate (EMS)–mutagenized population of *glk1 glk2*, leveraging an unbiased approach and identified an independent, extragenic regulator associated with enhanced chloroplast development, the *suppressor of glks 2* (*sgl2*). Our results show that loss of function of *SGL2* enhances chlorophyll content and photosynthetic gene expression in the near-absence of GLK factors. Mesophyll cells (M) are the primary contributors to photosynthesis, housing most of the chloroplasts in plants (Pyke, 2012; Tsukaya, 2013). Another main cell type, bundle sheath cells (BS), encircling vascular strands, contain fewer chloroplasts, constituting about 15% of photosynthetic cells in a dicotyledonous plant like *Arabidopsis* (Kinsman and Pyke, 1998). Investigation of the enhanced greening phenotype in *sgl2 glk1 glk2* showed improved cellular chloroplast compartment and thylakoid stacking. We show through bulk-segregant whole-genome sequencing analysis that *sgl2* carries a single nucleotide change resulting in a premature stop codon in the locus encoding *AUXIN RESPONSE FACTOR 2* (*ARF2*) and confirm the gene identity through complementation. The role of *arf2^sgl2^* is further validated by the rescue of *long hypocotyl 1* (*hy1*), a mutant severely defective in early chloroplast development.

Moreover, the overexpression of *GROWTH-REGULATING FACTOR 5* (*GRF5*), which controls cell and early plastid division and is negatively regulated by ARF2 (Beltramino et al., 2021), also mimicked the enhancement of chloroplast development in *glk1 glk2* to the same extent as *arf2^sgl2^.* However, our results showed that *glk1 glk2* plants retain a degree of *GLK1* expression, which was further enhanced in the *arf2^sgl2^ glk1 glk2* background. We also noted that the suppression phenotype of *arf2^sgl2^* was retained but reduced in the presence of null mutations of *glk1* and *glk2*, making KO alleles of *glk1* and *glk2* partially epistatic to *arf2^sgl2^*. We conclude that GRF5 regulates early stages of chloroplast development and is part of a gene circuit which acts in part ahead of, and in part through, the GLK1 factor, whether directly or indirectly. This gene circuit may open avenues to enhance photosynthesis in crops.

## Materials and methods

### Plant material, growth conditions and mutant genotyping

*Arabidopsis thaliana* wild-type ecotype Columbia (Col-0), *glk1 glk2* double mutant (Fitter et al., 2002), *35S:GLK1* (a gift from Prof. Jane Langdale, University of Oxford), *arf2-7* (SALK 24601), a presumed KO, *35S:GRF5* (a gift from Prof. Gorou Horiguchi, Rikkyo University) and *glk1*-2 *glk2*-2, CRISPR-generated (Han et al. 2024, a gift of Prof. Rob Larkin, Wuhan Agricultural University) were used. All mutants used in this study were in Col-0 background.

Sterilised seeds were stratified and germinated *in vitro* on MS media with 1% sucrose, under continuous white light, at a fluence rate of 100 µmol m^−2^ s^−1^, as previously described (Loudya et al., 2020). Plants transplanted on soil were grown at 16h photoperiod and a constant temperature of 21°C and fluorescent white light (colour 840) at 180 μmol m^−2^ s^−1^.

The T-DNA insertions lines, the point mutation and CRISPR-deletion mutations were genotyped based on PCR or PCR and restriction digestion followed by gel electrophoresis (Neff et al., 1998). PCR primers and relevant restriction enzymes are listed in Supplemental Tables S2-3.

### Chemical mutagenesis and isolation of *sgl2*

The *glk1 glk2* seeds (approximately 10,000) were chemically mutagenized using ethyl methane sulphonate for 4 hours as previously described (López-Juez and Hills, 2011). Healthy M1 plants grown in pools (over 50 per pool) produced M2 seeds including homozygous mutants.

Screening of germinated M2 seedlings showed a putative suppressor with enhanced greening phenotype, named as *sgl2*. The greening phenotype was confirmed in subsequent generations and by genotyping for the initial *glk1 glk2* mutations.

### Chlorophyll and protochlorophyllide measurement

Chlorophyll and protochlorophyllide pigments were extracted using light-grown or dark-grown seedlings or mature leaf samples in dimethylformamide. Pigments were quantified through spectrophotometry or spectrofluorimetry, following previously established methods (López-Juez et al., 1998; Vinti et al., 2000).

### Mapping by sequencing

Backcrossed *sgl2 glk1 glk2* plants with the un-mutagenised *glk1 glk2* parent resulted in F1 progeny showing an intermediate phenotype, mild green, between both the parents indicating a semi-dominant trait. Tissue samples collected from the reappeared suppressor mutants (over 120 plants) in the F2 generation and *glk1 glk2* plants were used for genomic DNA isolation. The whole genome sequence (Illumina) was analysed to identify the EMS induced polymorphisms using Easymap tool as previously described (Lup et al., 2023). SNPs with high allele frequency and altered amino acids identified putative candidates (Supplemental Table S1) for further analysis.

### Total proteins extraction and immunoblotting

Protein extraction was performed using cold acetone and urea following a previously described protocol (López-Juez and Hughes, 1995). Total proteins were transferred to a nitrocellulose membrane, blocked with 5% Blotto (Bio-Rad) or Intercept buffer (LI-COR), then incubated overnight with primary antibody in Blotto or Intercept buffer at 4°C. Immunoblotting was performed with primary antibodies (Agrisera): PsbO (AS06 142-33), LHCB1 (AS01 004) and HRP-conjugated goat-antirabbit secondary (AS09 607) or IRDye 800CW Goat anti-Rabbit IgG secondary antibody as previously described (Loudya et al., 2022). Proteins (Supplemental Table S4) were detected and imaged with an ChemiDoc (Bio-Rad) imaging system or detected and quantified using Odyssey (LI-COR) DLx Imaging System and Studio Lite Version 5.2 software.

### Vector construction and complementation

The *pARF2:ARF2* sequence was cloned using the Gateway® Technology. Genomic DNA was extracted from 5-day-old seedlings (Macherey-Nagel, Nucleospin^®^ Plant II DNA) and the *pARF2:ARF2* sequence was PCR-amplified using long adapter Gateway primers with gene specific primers which cover the presumed 2 kb promoter and 3’UTR of the gene (Supplemental Table S5). The PCR product was purified using the QIAprep Spin Miniprep kit (Qiagen). The pure amplicons with attB sites were initially recombined with the pDONR201™ entry vector and later into the destination vector pKGW (which carries attR sites). The final destination vector was transformed into *Agrobacterium* and through floral dipping into plants.

### Cell cycle analysis

Arabidopsis WT and mutant flower buds were analyzed for cell cycle stages (G1, S, G2+M) using flow cytometry as previously described (Loudya et al., 2021). Flower buds (10-15) were placed in a Petri dish with 100 µl Partec cystain UV precise P lysis buffer (Sysmex-Partec) and finely chopped with fine razor (Wilkinson) blades. The isolated nuclei were stained with 1 ml Partec cystain UV precise P Solution 2 DNA-binding DAPI. The solution, containing the chopped sample, was transferred to a glass flow cytometer cuvette, passed through a 20-30 µm filter and run through the flow cytometer (Sysmex CyFlow® Space, Sysmex). The DNA content (2C or 4C) was assessed by analyzing at least 10,000 nuclei per sample.

### Analysis of rosette growth

The 5-day-old seedlings grown on MS plates were transferred to soil trays in rows of 10, with mutant genotypes and wild-type controls randomly distributed. The soil composition, water treatment, temperature, and light intensity were maintained constant throughout the experiment. Images of rosettes were used to measure the area using the Fiji image analysis software (https://imagej.net/Fiji).

### DIC live microscopy and electron microscopy

Arabidopsis leaf samples were fixed in glutaraldehyde (3.5%) and Tween 20 solution, and incubated in the dark for 30 minutes. The samples were then vacuum infiltrated twice and incubated with EDTA at 65°C for 2 hours as previously described (Loudya et al., 2020). Leaf fragments were cut and teased into fine threads and mounted in a glycerol drop. Live quantitation was performed on Nikon Optiphot-2 microscope using NIS-Elements AR 2.30 software. Total number and area of the chloroplasts was measured in different planes.

For the TEM analysis, the mid regions (without mid vein) of young rosette leaves (30 day-old) were cut into ∼2-3 mm square sections and immediately dipped in a vial which contained about 200 µl of fixative (3% glutaraldehyde and 4% formaldehyde in a 0.1M Piperazine-N-N’ bis 2-ethanol sulfonic acid (PIPES) buffer). After embedding in resin blocks, the samples were sliced to make the ultra-thin and thick section to analyse chloroplast ultra structure and cell anatomy (Southampton Biomedical Imaging Unit) as described previously (López-Juez et al., 1998). The thylakoid quantitation parameters are described in Supplemental Table S6. For quantitation of thylakoids, grana stacks with a minimum of three lamellae were considered, and mean thickness was calculated by measuring up to 10 stacks per chloroplast section.

### RNA extraction, cDNA synthesis and quantitation of transcripts

RNA extraction was performed by following the Nucleospin^®^ Plant RNA (Macherey-Nagel) kit, following the manufacturer’s instructions, and including rDNase enzyme (RNase-free) use. Quality of RNA samples was determined by gel electrophoresis and RNA reverse transcribed using QuantiTect® Reverse Transcription (RT) kit (Qiagen). The cDNA was diluted to 10-fold prior to using quantitative PCR (Rotor-Gene Q real-time PCR cycler, Qiagen) and SyGreen Mix Lo-ROX (PCRBiosystems) mix. Primer pairs for RT-qPCR are listed in Supplemental Table S7.

### Genome copy number

Chloroplast genome (cpDNA) copy number was determined using qPCR with standard curve analysis as described previously (Loudya et al., 2020). Genomic sequences from three targets of the chloroplast genome (large single-copy region, small single-copy region, and inverted repeat region) and two nuclear (gDNA) genes were quantified along with standards of known concentration. The ratio of cpDNA:gDNA was calculated to quantify the absolute copies of cpDNA per haploid genome.

## Results

### Mutations in *GLK1* and *GLK2* severely affect chloroplast development yet allow sufficient plant growth

GLK1 and GLK2 TFs, identified more than three decades ago, are known as master regulators of chloroplast development. Mutations in these two factors lead to a pale green phenotype with small chloroplasts carrying perturbed thylakoid stacking in mesophyll cells (Fitter et al., 2002; Wang et al., 2013) (Figure 1A). However, the plants complete their life cycle without any severe damage. This demonstrates the existence of other regulators of chloroplast development which allow chlorophyll biosynthesis and thylakoid development in the absence of GLKs. We employed an in-depth EMS mutagenesis screen in *glk1 glk2* to identify such regulators (Figure 1B). Rigorous screening of a mutagenized M2 population obtained a seedling with an enhanced greening phenotype, referred as a putative suppressor of *glk1 glk2*.

**Figure 1.**
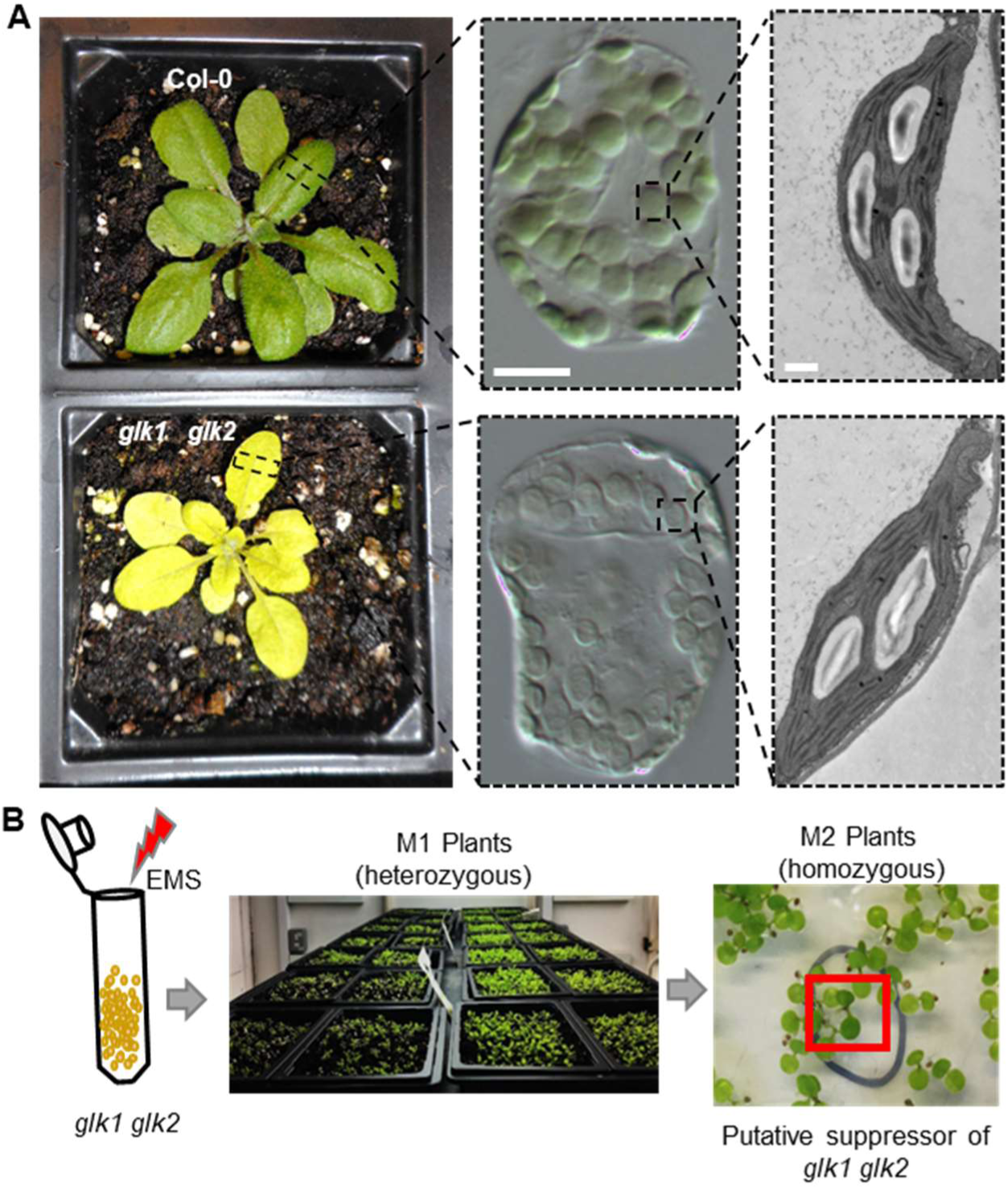
Morphological, cellular phenotype and schematic of genetic screen in *glk1 glk2*. (A) Phenotype of 20-day-old *glk1 glk2* depicting pale green phenotype along with wild type plant. Differential Interference Contrast (DIC) and electron micrographs showing chloroplasts in mesophyll cell and their ultrastructure. Scale bar 20 µm and 1 µm respectively. (B) Illustration of EMS mutagenesis in *glk1 glk2* and screening of suppressor plants in M2 population, red box depicting identified suppressor of *glk1 glk2*.

### Forward genetic screen in *glk1 glk2* identifies a suppressor with enhanced greening

The greening phenotype of the identified putative suppressor was confirmed in the next generation and the mutant named ***s****uppressor of **gl**ks 2* (***sgl2***). Plants of *sgl2 glk1 glk2* exhibited a substantial phenotypic suppression of *glk1 glk2* (Figure 2A). Analysis of *sgl2 glk1 glk2* showed a doubling in total chlorophyll content compared to *glk1 glk2* (Figure 2B). The *glk1 glk2* mutant also contained reduced levels of photosynthesis-related proteins and transcripts. *sgl2 glk1 glk2* displayed increased transcript levels of photosynthesis-associated nuclear (PhAN) genes (Figure 2C). Quantitation of photosynthetic component proteins, specifically the photosystem II (PSII) antenna (LHCB1) and a nucleus-encoded protein of the PSII reaction centre (PsbO), which were reduced in *glk1 glk2*, showed a partial rescue by *sgl2*: the reduction of PsbO protein level was no-longer statistically significant (Figure 2, D-F). Backcrossing *sgl2 glk1 glk2* into the *glk1 glk2* parent resulted in an F1 with an intermediate phenotype between *glk1 glk2* and *sgl2 glk1 glk2* (Supplemental Figure S1, A and B) indicating that *sgl2* is a semidominant mutation. Overall, the phenotypic analysis (Figure 2, A-F) suggests that the *sgl2* mutation rescues chlorophyll biosynthesis and to some extent the accumulation of photosynthetic proteins in the *glk1 glk2* mutant.

**Figure 2.**
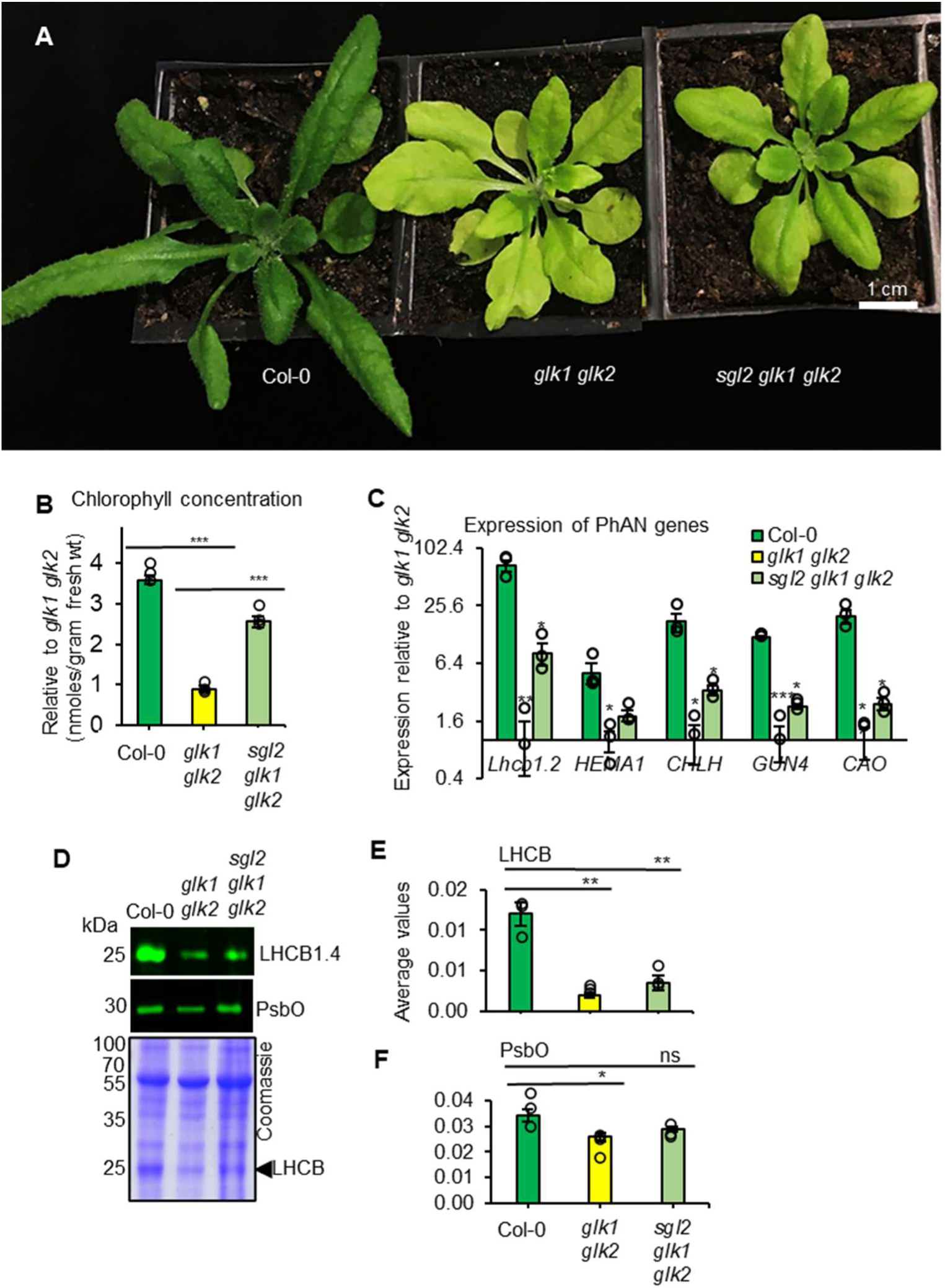
Morphological phenotype of the suppressor and its suppression parameters.(A) Image of potential suppressor *sgl2 glk1 glk2*, compared to the controls *glk1 glk2* and Col-0 grown for 20 days under 16-hour photoperiods, 180 µmol/m²/s light at 21°C. Scale bar 1 cm. (B) Chlorophyll quantitation per rosette leaf gram fresh weight shown relative to *glk1 glk2*. (C) Transcript levels of PhAN genes in *sgl2 glk1 glk2*, *glk1 glk2* and Col-0, expressed on a log₂ scale relative to *glk1 glk2*. (D) 20 µg of total protein from the indicated genotypes was run on a 10% polyacrylamide gel alongside a ladder (PageRuler™ Plus Prestained Protein Ladder). The membrane with protein was incubated for 1 hour using anti-LHCB and anti-PsbO primary and fluorescent secondary antibodies. Protein detection was performed using the Odyssey (Li-Cor) DLx Infrared imaging system. Equal loading of protein from the same samples is shown in the Coomassie-stained gel. Protein band intensity was measured using Image Studio Lite Ver 5.2 by comparing selected Coomassie-stained protein bands (65 to 120 kDa) above RuBisCO large subunit (ca. 55 kDa) for the respective sample. Arrowhead indicates LHCB protein bands. Quantitation of (E) LHCB and (F) PsbO levels in indicated genotype. All the experiments were performed in 20-day old rosette leaves. The error bars represent SEM values (n=3, independent immunoblots). Asterisk with the Student’s t-test P values indicates significant difference between mutant and WT ≤0.05 (*), ≤0.01 (**) and ≤0.001 (***).

### Identification of the *sgl2* locus

Mapping by sequencing was carried out by backcrossing (BC) *sgl2 glk1 glk2* with *glk1 glk2*. In the F2 generation, green plants resembling the suppressor mutant phenotype were selected. Paired-end next-generation sequencing (NGS) reads of the BC1F2 *sgl2 glk1 glk2* genome were compared with those of the non-mutagenized *glk1 glk2* (Supplemental Figure 2A) using the Easymap analysis (Lup et al., 2023). Single nucleotide polymorphisms (SNPs) present in chromosomes 1-4 showed random distribution, whereas a clear pattern was observed in chromosome 5 (Figure 3A and Supplemental Table S1) including a C to T nucleotide change (position 24913807) in AT5G62000, which introduced a premature stop codon in place of a glutamine at the 628^th^ position of the amino acid sequence. This point mutation on exon 12, indicated in the gene structure (Figure 3B) was genotyped and confirmed through a derived Cleaved Amplified Polymorphic Sequence (dCAPS) assay (Figure 3C).

**Figure 3.**
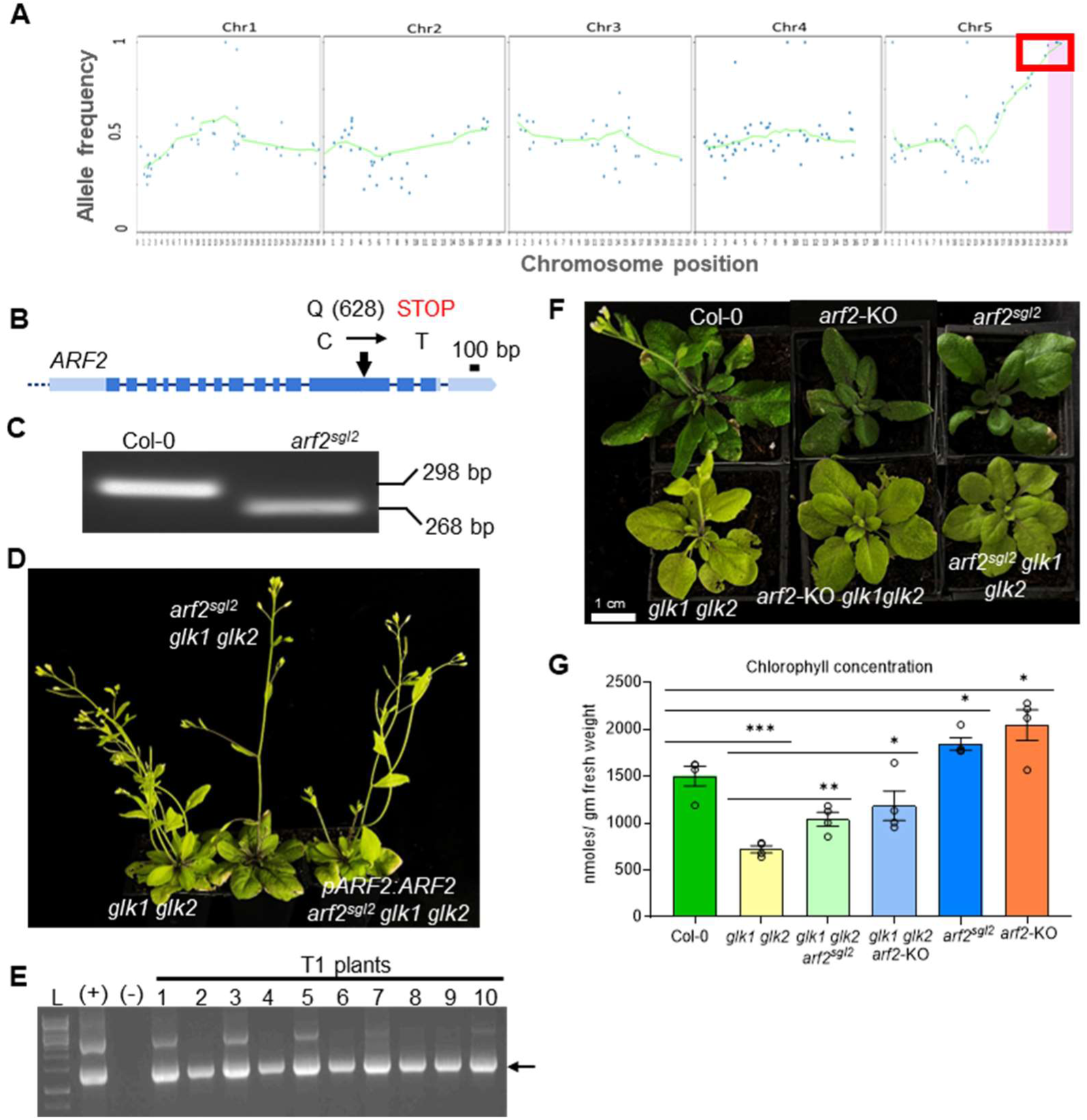
Identification of the *sgl2* locus through mapping by sequencing and confirmation of s*g*l2 identity. (A) Position of EMS-generated polymorphisms in different chromosomes (location in Mb on x-axis) and their allelic frequency (y-axis). (B) Gene structure of the likely candidate and location of nucleotide change, C to T transition, which leads to premature STOP codon generation. (C) Mutant genotyping through dCAPS assay. (D) Complementation of suppressor mutation functional copy of *ARF2* including its presumed full-length promoter. Transformed *arf2^sgl2^ glk1 glk2* plant showing pale phenotype resembling *glk1 glk2* in T2 generation. (E) Genotyping of transformants of *arf2^sgl2^glk1 glk2* in the presence of positive and negative control. (F) Phenotype of 20-day-old *arf2-KO glk1 glk2* and *arf2^sgl2^ glk1 glk2* with their controls. (G)) Chlorophyll concentration in *arf2 glk1 glk2*, *arf2^sgl2^ glk1 glk2* with their controls. A significant difference is presented by an asterisk with the Student’s t-test P value ≤0.05 (*), ≤0.01 (**) and ≤0.001 (***). Error bars represent SEM values (n≥3).

*sgl2 glk1 glk2* plants were transformed with *Agrobacterium* carrying a vector containing genomic DNA of *ARF2* with its presumed complete native promoter (1985 bp upstream of *ARF2*). The suppressor mutant exhibits a semi-dominant phenotype; remarkably, 10 selected T1 transformants which were confirmed for the presence of the transgene (Figure 3E) showed an intermediate phenotype. Their T2 generation showed plants which fully phenocopied *glk1 glk2* (Figure 3, D and E), demonstrating that the functional copy of *ARF2* complements and rescues (reverts to) the *glk1 glk2* phenotype. Hereafter the *sgl2* mutation, and mutant plant carrying it, will be referred as *arf2^sgl2^*.

To further support this conclusion, a separate *arf2*-KO allele, carrying a T-DNA insertion in the *ARF2* gene (Supplemental Figure 2, B and C), was crossed with *glk1 glk2*. The homozygous triple *arf2*-KO *glk1 glk2* mutant showed suppression of the pale *glk1 glk2* phenotype, completely mimicking the greening of *arf2^sgl2^glk1 glk2* (Figure 3F) and accumulated similar levels of chlorophyll (Figure 3G). Following backcrossing with the wild type we identified the *arf2^sgl2^* single mutant. Both single mutants, *arf2^sgl2^* and *arf2*-KO, showed enhanced greening compared to wild-type at the seedling (Supplemental Figure 2, D and E) and mature stage (Figure 3G). The results demonstrate that the *sgl2* mutation in *ARF2* is indeed responsible for the suppression of *glk1 glk2*.

### *arf2^sgl2^* rescues the chloroplast defects in *glk1 glk2*

Loss of function of GLK factors in maize (Langdale and Kidner, 1994), Arabidopsis (Fitter et al., 2002), rice (Wang et al., 2013), and *Setaria* (Lambret-Frotte et al., 2024) affects the chloroplast content (coverage) of cells significantly. Fitter et al. (2002) initially showed that *glk1 glk2* cells have reduced chloroplast size when compared with those of developmentally similar rosette leaves of wild-type. We performed live chloroplast quantitation in mesophyll cells, which predominantly represent the leaf chloroplast population (Pyke, 2011; Loudya et al., 2020). We hypothesized, considering the impact on greening, that *arf2^sgl2^* might improve the chloroplast compartment in *glk1 glk2* and quantified the chloroplast number, size and cellular content (cellular chloroplast index) in *arf2^sgl2^ glk1 glk2*. *glk1 glk2* carried much smaller chloroplasts, a mean area reduction to around 60%, in mesophyll cells, and this was fully rescued by *arf2^sgl2^ glk1 glk2* (Figure 4, A and B). While GLK factors influence plastid development and differentiation, they do not impact plastid division (Fitter et al., 2002), and indeed the chloroplast count per cell remained normal in double mutants. Surprisingly, the *arf2^sgl2^ glk1 glk2* exhibited a significant increase in chloroplast density and total chloroplast area per cell, even higher than those of the wild-type (Figure 4, C and D).

**Figure 4.**
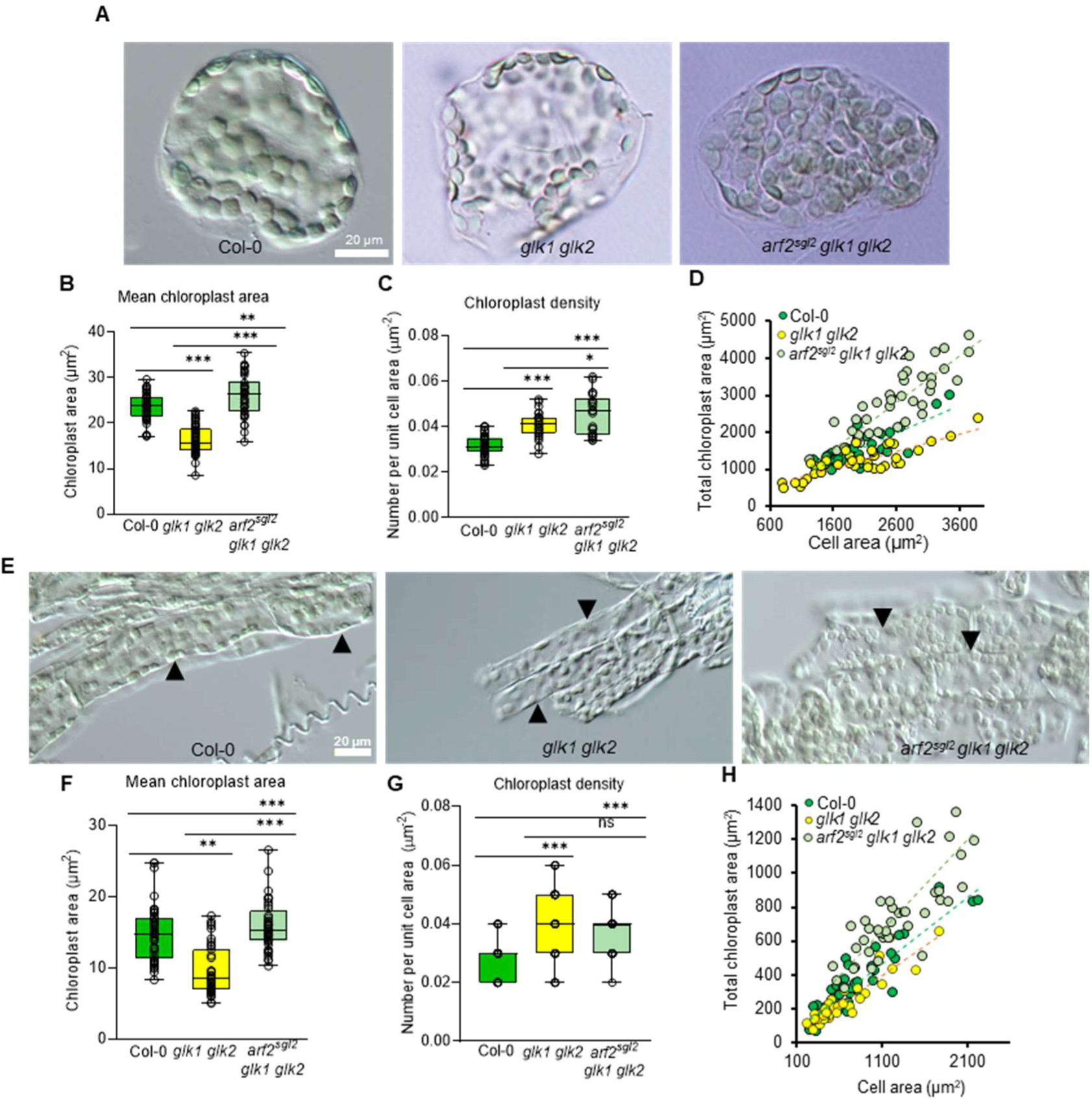
Chloroplast cellular parameters of *arf2^sgl2^ glk1 glk2*. (A) Mesophyll cells, obtained from fixed leaves of the genotypes mentioned. (B) Mean area of individual chloroplasts.(C) Chloroplast density, number per unit area, (D) Total chloroplast area represented against each cell’s plan area. (E) Chloroplasts in bundle sheath cells of *arf2^sgl2^ glk1 glk2* and controls. Individual bundle sheath cells are highlighted by black arrowheads.(F-H) Individual chloroplast area (F), chloroplast density (G) and total chloroplast area against cell area of bundle sheath cells (H). Cells obtained from 20-day-old rosette leaves, 40 cells quantified per genotype. The error bars represent ± SEM (n = 3 independent leaves). The asterisk symbol represents a significant difference between the mutants and their controls as determined by individual Student’s t-test P-values ≤0.05 (*), ≤0.01 (**) and ≤0.001 (***).

In general bundle sheath cells (BS), a second leaf cell type, contain smaller chloroplasts in C3 plants (Kinsman and Pyke, 1998; Wang et al., 2013). Mutations in *GLKs* make those chloroplasts even smaller (Wang et al., 2013). The BS in *arf2^sgl2^ glk1 glk2* showed fully rescued chloroplast size and double the chloroplast numbers when compared to *glk1 glk2* but surprisingly also showed increased cellular chloroplast index even compared to wild-type cells (Figure 4, E-H). However the number of copies of the chloroplast genome of the leaf overall, per haploid nuclear genome, in *arf2^sgl2^ glk1 glk2* and *glk1 glk2*, remained similar to that of wild type (Supplemental Figure 3 K, L). The significant increase in the total chloroplast area in M and BS cells relative to cell area was further evidenced in the *arf2^sgl2^* single mutant (Supplemental Figure 3, A-J) suggesting *ARF2* regulates the overall chloroplast number (per unit area) and size in photosynthetic cells, even in the presence of GLK factors.

### *arf2^sgl2^* reestablishes thylakoid development in *glk1 glk2*

Loss of function in GLKs results in reduced chloroplast compartment and thylakoid development in various plant species (Langdale and Kidner, 1994; Fitter et al., 2002; Wang et al., 2013; Frangedakis et al., 2024). We carried out transmission electron microscopy in the suppressed *glk1 glk2* and *arf2^sgl2^* lines using rosette leaf tissue, to analyse chloroplast ultrastructure. This analysis included the number of grana stacks (with three or more lamellae), mean thickness of the stacks, total grana thickness per chloroplast section (number of grana stacks × mean thickness of thylakoid stack), and "grana load" (total grana thickness per unit chloroplast area). The ultrastructure of *arf2^sgl2^ glk1 glk2* and *arf2^sgl2^* lines displayed improved thylakoid development (Figure 5, A-C). The larger chloroplasts in the suppressed *glk1 glk2* lines contained more than double the number of thylakoids (Figure 5, D) and much thicker grana compared to *glk1 glk2* (Figure 5, E). In the single *arf2^sgl2^*mutant, overall thylakoid development (thylakoid number and grana thickness) was enhanced (both by over a third) even compared to WT. Consequently, total grana thickness and grana load were significantly enhanced as well (Figure 5, F and G). Taken together, transmission electron microscopy analysis of *arf2^sgl2^* showed a clear positive impact on thylakoid and overall chloroplast development, correcting the *glk1 glk2* mutant but also occurring in the presence of GLK factors.

**Figure 5.**
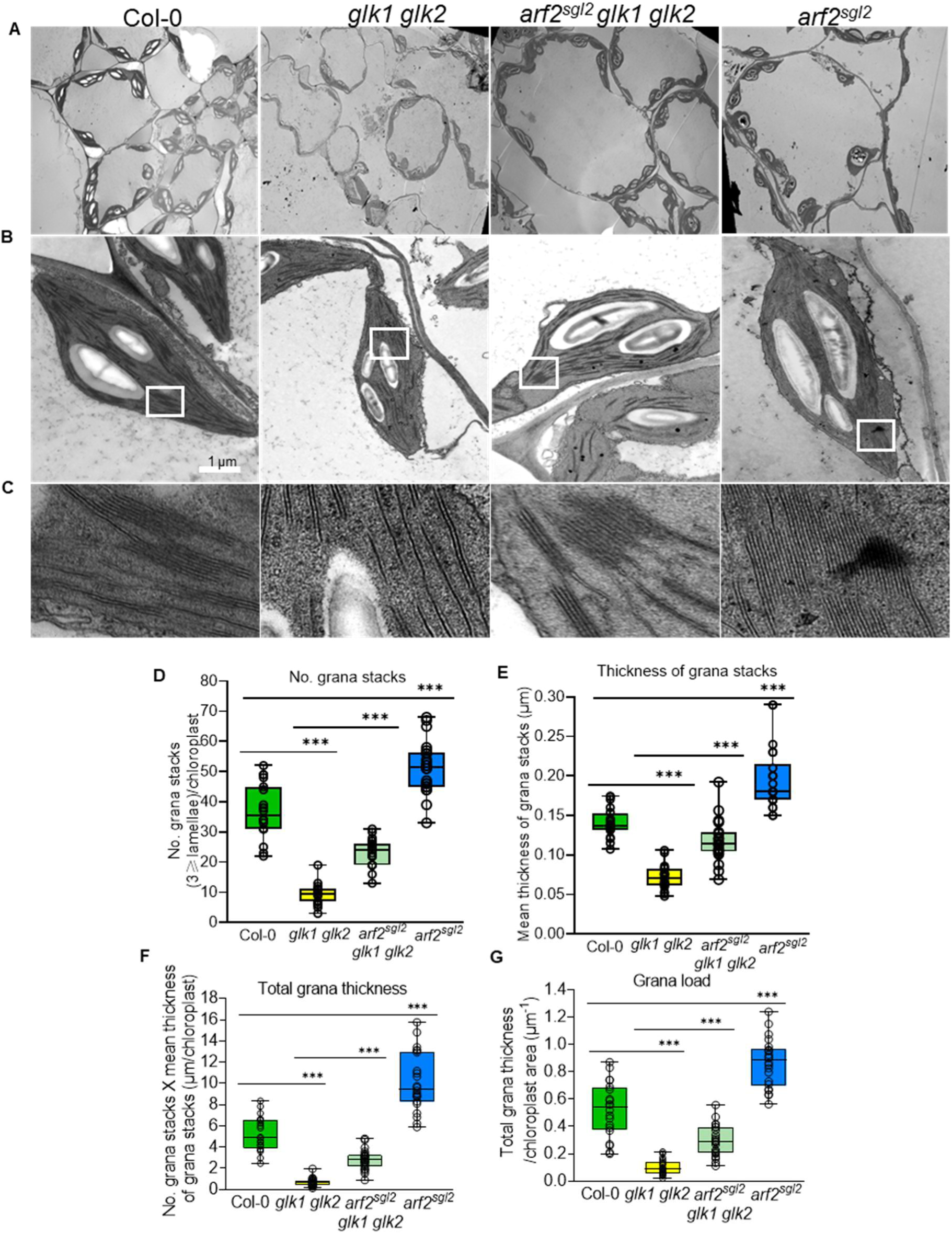
Chloroplasts ultrastructure and its analysis. (A) Mesophyll chloroplasts of both the mutant and wild-type (Col-0) captured at 1250x magnification, and (B) at 11500x magnification. (C) The magnified region of the thylakoids from chloroplasts in (B) is indicated by a white box. (D, E, F, and G) Quantitative analysis includes the number of grana, mean thickness, total grana thickness, and grana load. Analysis was conducted using 30-day-old young rosette leaves (middle section of the leaves). Error bars represent ± SEM (n = 2 replicates; number of chloroplasts ≥ 20). White structures within the chloroplasts are starch granules. A significant difference between the mutant and WT is indicated by Student’s t-test P-values ≤0.001 (**).

Additionally, the level of thylakoid-localised chlorophyll binding protein (LHCB1) appeared slightly increased in *arf2^sgl2^* and *arf2-*KO (Supplemental Figure 4A). Disruption of GLKs function also impairs development of dark-grown etioplasts (Waters et al., 2009), with protochlorophyllide (a precursor to chlorophyll) content being significantly reduced. The protochlorophyllide content of dark-grown *arf2^sgl2^* seedlings showed a mild but significant increase compared to the wild type (Supplemental Figure 4, B and C). Investigation of different parameters associated with chloroplast development therefore suggests that the loss of function of *ARF2* further enhances thylakoid biogenesis and plastid development in both dark and light conditions.

### *arf2^sgl2^* flowers late, but the correction of the chloroplast defect is independent of flowering

When grown under long day conditions, the *arf2^sgl2^* plants displayed larger total rosette area compared to the WT, with an increased average number of rosette leaves per plant from 13 (WT) to 19 (*arf2^sgl2^*), and with a consistent increase in chlorophyll content (Figure 6, A-D). The *arf2^sgl2^ glk1 glk2* plants showed delayed flowering, by 10 days (Supplemental Figure 5, A and B) under long day conditions, with much larger plants resulting. While a delay in flowering has been observed following the overexpression of positive chloroplast regulators - GLKs, GNL, and GRF5 (Waters et al., 2008; Richter et al., 2010; Vercruyssen et al., 2015), this is a greater delay. Analysis of leaf anatomy revealed a severe reduction of cell layers in *glk1 glk2*, a phenotype which is fully rescued in the *arf2^sgl2^ glk1 glk2* (Supplemental Figure 5C). We asked whether the enhanced greening is due to the delayed flowering response or to enhanced chloroplast development, by growing plants under short-day conditions (SD) in which flowering is greatly delayed in all genotypes. *arf2^sgl2^* exhibited a slightly reduced rosette size relative to its control, while *arf2^sgl2^ glk1 glk2* was unaltered, but both mutants showed an increased chlorophyll concentration in SD compared with their controls (Figure 6, E-G). *arf2^sgl2^ glk1 glk2* and *arf2^sgl2^* plants are much larger than their controls in long days, and studies have shown a correlation between plant size and ploidy: in Arabidopsis an increase in somatic ploidy to 4n and, to some extent, 8n is associated with increased plant size and biomass (Corneillie et al., 2019). This, however, was not the cause of the increased size of our mutant: flow cytometry analysis showed unaltered ploidy levels in *arf2^sgl2^* compared to wild type (Supplemental Figure 5, D and E). Furthermore, the *arf2^sgl2^* plants also showed increased seed size (Supplemental Figure 5, F and G). These observations show that the increase in greening in *arf2^sgl2^* plants is not simply caused by the delay in flowering, and that the overall increase in plant growth, which causes even enlarged seeds, is not due to altered ploidy levels.

**Figure 6.**
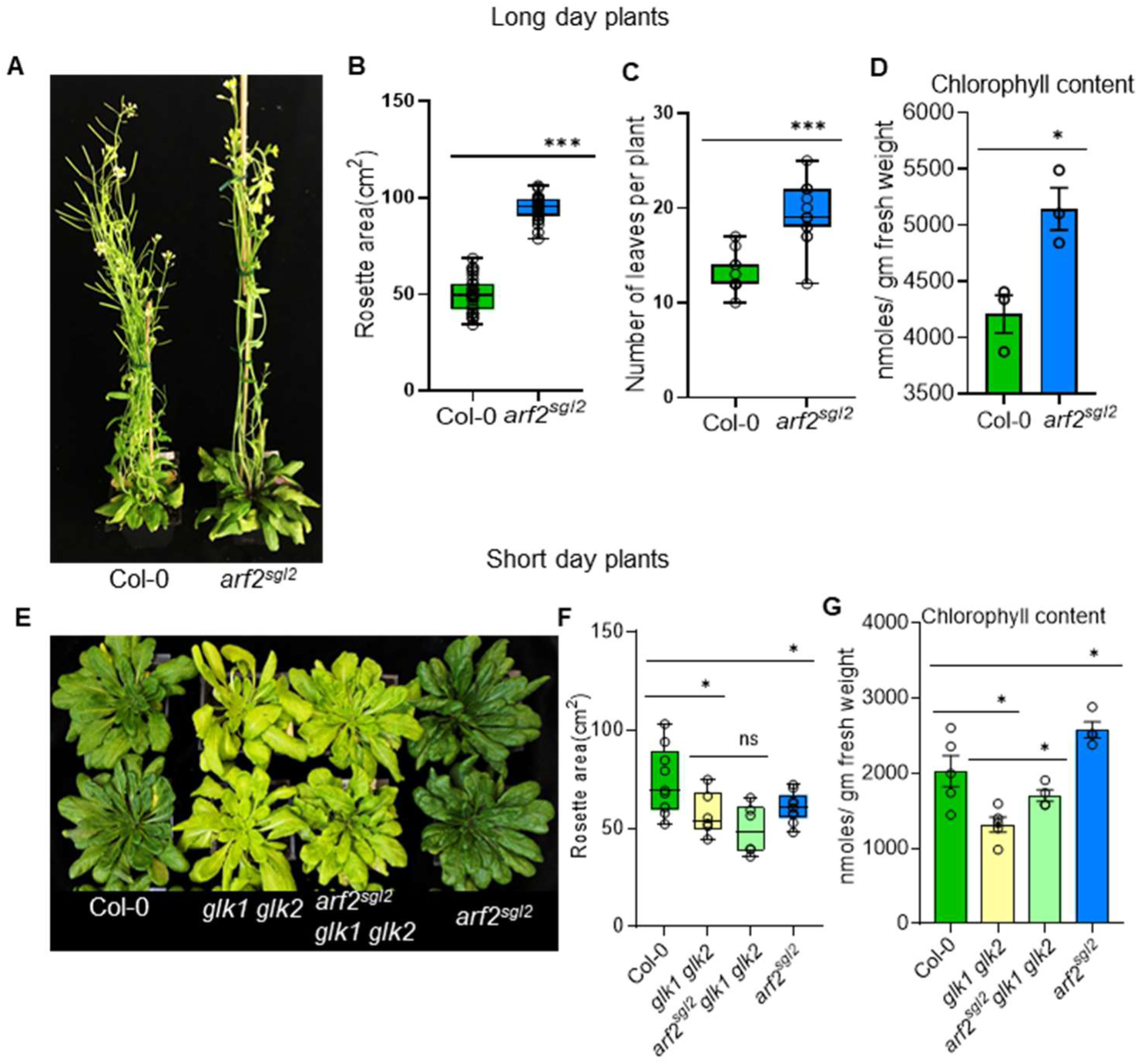
Description of rosette growth and greening in long-day and short-day conditions. (A) Plant phenotype of the 35-day old mutant and wild type in long day conditions. (B) Area of the rosette of leaves. (C) Rosette leaf number per plant. (D) Chlorophyll levels per unit leaf weight. (E) Phenotype of the 45-day old mutant and wild type in short day conditions. (F) Average rosette area per plant and (G) chlorophyll concentration. The asterisk symbol represents a significant difference between the mutants and their controls using Student’s t-test P-values ≤0.05 (*) and ≤0.001 (***).Error bars represent ± SEM (n ≥ 9 plants).

### Genetic interaction supports the impact of *arf2* enhancing chloroplast development

GLK transcription factors (TFs) activate photosynthesis-related gene expression after light induction (Fitter et al., 2002; Waters et al., 2009; Wang et al., 2013). Between the two TFs, the transcript levels of *GLK2* show a strong response to early light induction that depends on phytochrome (Tepperman et al., 2006). Photosynthesis is influenced by red light-perceiving phytochromes that function upstream of GLK factors. The production of active phytochrome formation requires phytochromobilin synthesis in the chloroplasts. The *long-hypocotyl1* (*hy1*-100) null mutant, defective in a phytochromobilin synthesis rate-limiting enzyme, HAEM OXYGENASE 1, shows greatly impaired light signalling, plant growth (Muramoto et al., 1999) and chloroplast development (Vinti et al., 2000). To investigate whether *arf2^sgl2^*suppresses chloroplast development defects upstream of regulation by GLK factors, we generated *arf2^sgl2^ hy1*-100 (Supplemental Figure 6 A, B). Loss of function of *ARF2* dramatically rescued the phenotype of *hy1*-100 before (Supplemental Figure 6 A, B) and after flowering, by increasing the number of leaves per plant and also their development (Figure 7, A-C) and greening (Figure 7D). The mesophyll cells of *hy1*-100 possess smaller chloroplasts which fail to fill the cellular space but much of the reduction in cellular chloroplast index was reverted in the *arf2^sgl2^ hy1*-100 plants (Figure 7, E-H). Our results highlight that *arf2^sgl2^* enhances chloroplast development and plant growth even in a phytochrome-deficient background.

**Figure 7.**
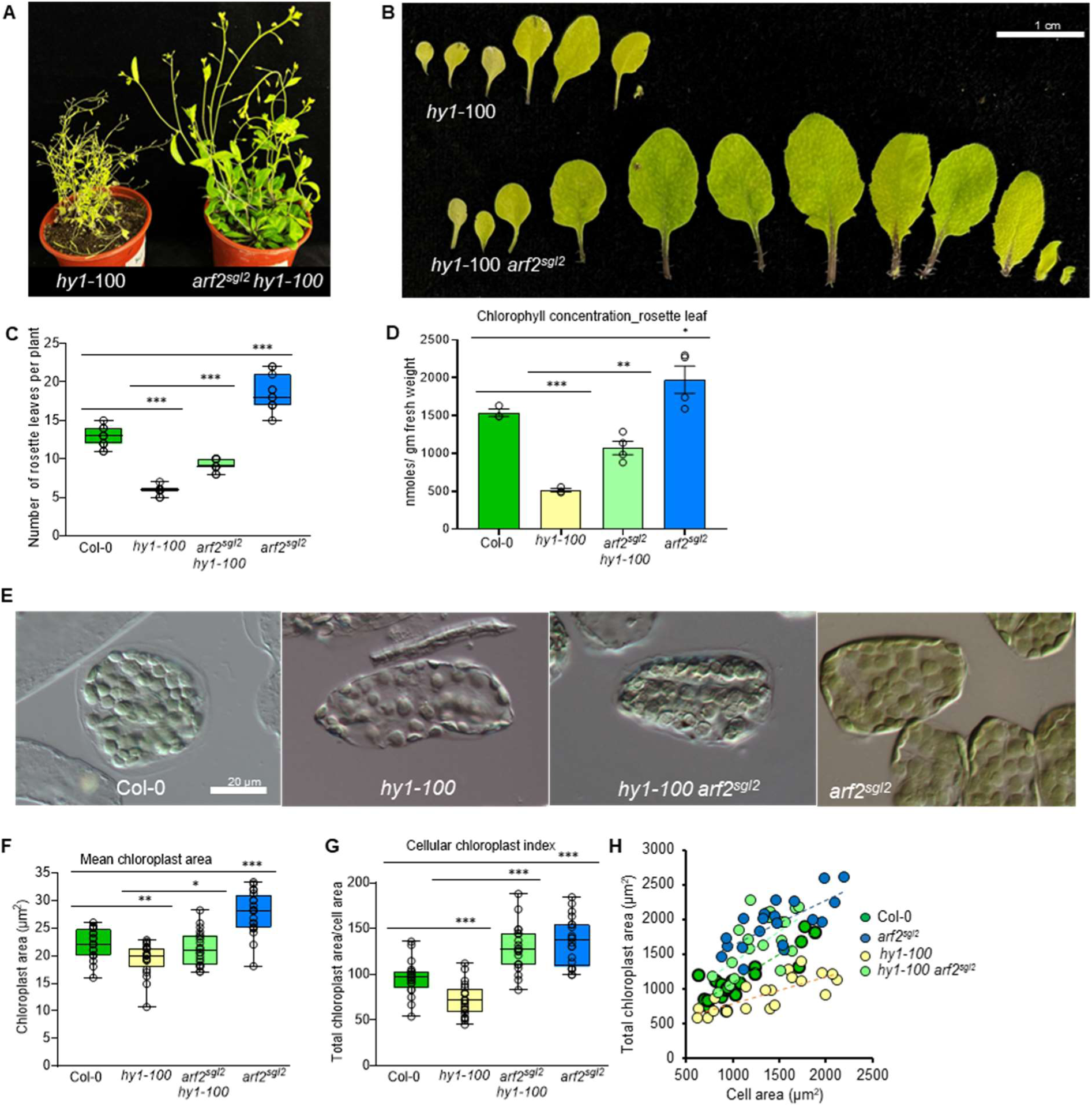
Genetic Interaction of *arf2^sgl2^* and the phytochrome-deficient *hy1*-100 mutant. (A) Suppression phenotype of *hy1 arf2^sgl2^* (B and C) Developmental phenotype and number of rosette leaves.(D) Chlorophyll content in 30-day-old rosette leaves. (E) Mesophyll cells of the genotypes indicated. (F-H) Chloroplast parameters. Error bars represent ± SEM (n = 3 independent leaf samples, number of cells = 20). Significant differences are indicated by Student’s t-test P-values ≤0.05 (*), ≤0.01 (**), and ≤0.001 (***).

### The action of *arf2^sgl2^* is partly dependent on expression of the *GLK1* gene

The *glk1 glk2* mutant carries T-DNA insertions in the proximal 5’ UTR of the transcription start site of *GLK1* and within the coding region of *GLK2* (Fitter et al., 2002). While RNA gel blots fail to detect any transcript of *GLK1* in this mutant, we sought to quantify *GLK1* and *GLK2* transcript levels in it and in the suppressed mutant by quantitative real-time PCR (qPCR) (Figure 8A). To our surprise, and consistent with observations of Quevedo et al. (2025), while the transcript of *GLK2* was essentially undetectable in the *glk1 glk2* and *arf2^sgl2^ glk1 glk2* mutants, that of *GLK1* was detectable, about 1/10 of wild type level in the *glk1 glk2* mutant, and this was doubled, to about 1/5 of wild type, in the triple mutant (Figure 8A).

**Figure 8.**
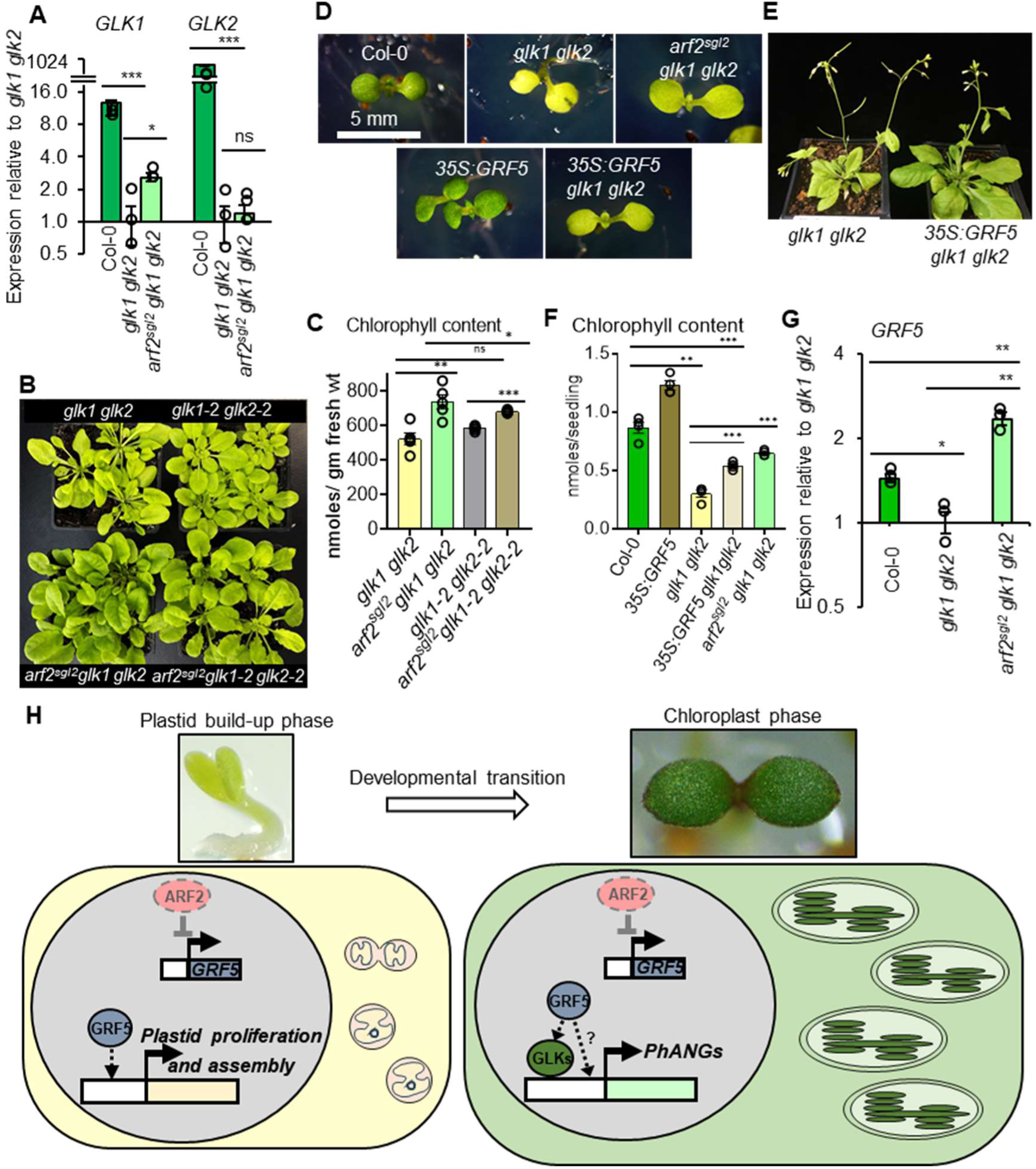
Suppression of the *glk1 glk2* phenotype through overexpression of *GRF5* and elevated *GLK1* expression. (A) Expression of *GLK1* and *GLK2* in *arf2^sgl2^ glk1 glk2* and its controls, revealing the likely knock-down nature of the *glk1* mutation in this *glk1* allele. (B) Partial suppression phenotype of the *arf2^sgl2^ glk1*-2 *glk2*-2, knock-out alleles. (C) Chlorophyll quantitation in the lines as labelled, double and triple mutants differing in the nature of the *glk1* and *glk2* alleles. (D and E) Suppression phenotype at seedling and flowering stage in *35S:GRF5 glk1 glk2*. (F) Accumulation of chlorophyll in 7-day-old seedlings. Error bars represent ± SEM (n ≥ 3). (G) Transcript levels of *GRF5* in fully expanded leaves of 20-day-old plants.(Error bars represent ± SEM (n = 3). (H) Working model illustrating that the ARF2-GRF5 module can effectively promote chloroplast development in early stages by promoting the early plastid proliferation and assembly phases (plastid stage). Removing ARF2 may prolong plastid division during early development while also enhancing the developmental retrograde signalling mechanisms which, via GLKs and yet to be determined factors, directly or indirectly, drives expression of chlorophyll biosynthesis genes during the greening phase. This combined effect enhances overall chloroplast development. Our findings demonstrate that the ARF2-GRF5 module can effectively promote chloroplast development in part independently of the GLKs, in part by boosting them, highlighting the hierarchical regulatory cascade of chloroplast biogenesis.

These results were consistent even in the *arf2^sgl2^ hy1-100* (Supplemental Figure 7 A). To assess the significance of this expression level, we used a recently available CRISPR-generated *glk* double mutant (Han et al., 2024), carrying alleles in which a section of the genomic coding region if missing for both genes, additionally causing frame shifts, resulting in null alleles. The *arf2^sgl2^ glk1*-2 *glk2*-2 mutants still showed suppression of the *glk1*-2 *glk2*-2 phenotype, but the degree of suppression was about halved, relative to that of the previous *glk* alleles (Figure 8B and C). This reveals that part of the suppression of the *glk* mutant phenotype by *arf2^sgl2^* is due to the enhancement of the very weak expression of *GLK1*, and consequently that ARF2 acts partly upstream of GLK factors, through a common chloroplast biogenesis gene-regulatory pathway, and partly in parallel to GLK factors.

### An ARF2-GRF5 module is sufficient to regulate chloroplast development

(Beltramino et al., 2021) demonstrated that ARF2 is a negative regulator of *GROWTH REGULATING FACTOR5* (*GRF5*). The over-expression of *GRF5* causes enhanced cell division and growth and, perhaps as a consequence of the enlarged cells, increased plastid numbers per cell, which overall enables increased leaf development and greening (Horiguchi et al., 2005; Vercruyssen et al., 2015). We asked whether the greening phenotype in *arf2^sgl2^ glk1 glk2* is achieved by the elevated expression of *GRF5*. We used a previously-characterised *35S:GRF5* line (Horiguchi et al., 2005) to generate a triple *35S:GRF5 glk1 glk2* mutant line (Figure 8D and Supplemental Figure 7, B-D).

Constitutive expression of *GRF5* suppressed the *glk1 glk2* phenotype. The enhanced chlorophyll concentration in *35S:GRF5 glk1 glk2* was apparent from the early seedling stage and became more pronounced later in development (Figure 8D-F). The triple lines also showed an overall boost in plant development as evidenced by larger leaves, similar to those of *arf2^sgl2^ glk1 glk2* (Figure 8E), while the increase in plant length in *arf2^sgl2^* was comparable to that of *35S:GRF5* (Supplemental Figure 7, E-H). *arf2^sgl2^* indeed caused elevated expression of *GRF5* in the *glk1 glk2* background, although the degree of overexpression was much greater in *35S:GRF5 glk1 glk2* (Figure 8G, Supplemental Figure 7C). Taken together, these analyses indicate that suppression in *arf2^sgl2^ glk1 glk2* is due to the enhanced expression of *GRF5*. Nevertheless, although both *35S:GRF5* and *arf2^sgl2^* cause an obvious greening phenotype and increased chloroplast index, the number of plastid genome (cpDNA) copies relative to their haploid nuclear genome remained similar to wild-type, implying that the increase in chloroplast number and size in these lines did not involve changes in cpDNA content (Supplemental Figure 7I).

## Discussion

The process of chloroplast biogenesis is required in vascular plants owing to the fact that leaf organs develop from pools of stem cells, initials, located at shoot apical meristems and with a very different phenotype to that of mature cells. Such stem cells are comparatively very small, with dense cytoplasms and minimal vacuoles, and have evolved to carry simplified chloroplast precursors, proplastids or “eoplasts” (Jarvis and López-Juez, 2013). Stem cells which will differentiate as mesophyll cells, those in the layer below the epidermis, enter a phase of extraordinarily active proliferation before their expansion and their reaching a fully differentiated, photosynthetically-active state in which they are filled with mature chloroplasts. This sequence of events, which is particularly apparent in cereal leaves, displayed at different points from the meristematic base to the mature leaf tip, has been described by classic studies as operating in stages (Leech et al., 1973; Mullet, 1993): plastids proliferate and establish housekeeping functions, before becoming green chloroplasts. Genome-enabled analyses have dramatically elaborated on those descriptions (Li et al., 2010; Wang et al., 2014; Chotewutmontri and Barkan, 2016; Loudya et al., 2021; Loudya et al., 2024), defining what can be best described as three phases: plastid proliferation (including the bulk of genome replication, coinciding but extending beyond cell proliferation), plastid assembly (in which the internal transcription machinery operates and that for internal translation and for external protein import is established), and greening phase, in which thylakoid membranes and photosynthetic complexes are themselves assembled.

One particularly highly-resolved proplastid-to-chloroplast development analysis using cellular, molecular and transcriptomics observations in the developing wheat leaf allowed visualisation of gene expression patterns along the developmental gradient (Loudya et al., 2021). The profile of *GLK* transcripts showed extended expression during the later, greening stages, as could have been assumed from the fact that *GLK*-dependent genes overwhelmingly include those for photosynthetic complexes or involved in light-harvesting pigment biosynthesis (Waters et al., 2009). This highlighted the need of earlier plastid biogenesis regulators active before these TFs (Wang et al., 2017; Loudya et al., 2021). It is particularly revealing to confirm that, in contrast to *GLKs*, the expression of the closest homologues of both *ARF2* and *GRF5* peaks during early development at the base of the developing wheat leaf (Supplemental Figure 7J).

Early in leaf development hormones play fundamental roles in leaf initiation and development (Braybrook and Kuhlemeier, 2010; Tsukaya, 2013). Auxin maxima determine the locations at which leaf primordia will be established. However it is the export of auxin, which is light-dependent, what will allow the establishment of a site of high cytokinin activity, accompanied by rapid cellular proliferation and cytoplasmic growth, i.e. translation capacity (Mohammed et al., 2018). This results in leaf primordium development. Cytokinin is a driver of both leaf primordium cell proliferation and greening (Cackett et al., 2022). Auxin export is necessary to allow leaf initiation in the light (Mohammed et al., 2018; Dóczi et al., 2019) because of the very well-established antagonism between auxin and cytokinin action (Kurepa et al., 2019). In this regard it is important to note that GRF5 is a target of cytokinin action (Horiguchi et al., 2005; Vercruyssen et al., 2015), while its repressor, ARF2 (Beltramino et al., 2021), is an auxin response factor.

Meanwhile, a close association and bi-directional control between leaf development and chloroplast biogenesis has been identified and documented (Andriankaja et al., 2012), for example by observing that preventing chloroplasts from progressing past the organelle proliferation stage delays exit of cells from the active cell cycle. The analysis of the plastid-to-chloroplast biogenesis sequence in the developing wheat leaf (Loudya et al., 2021) revealed that states of early plastid proliferation and assembly exhibit features which we and others had previously observed as phenotypes of chloroplast development mutants (Loudya et al., 2020; Kendrick et al., 2022). This led to a model in which defects in those mutants could be explained by failure to transition through consecutive stages in chloroplast biogenesis.

Stages of proliferation, assembly and greening follow on each other in a common developmental trajectory, subject to successful progress through checkpoints resulting from the occurrence of plastid-to-nucleus signalling events (Loudya et al., 2024). Such a model explains many experimental observations on both organelle and leaf development, in Arabidopsis, maize and wheat (Loudya et al., 2024). The data presented here support and expand on such a model: as part of this trajectory, cell and plastid proliferation stages may be dependent on hormone-target genes, and successful progress through this phase may contribute to passage through to the greening phase by promotion of the greening drivers.

Early GRF5 action, which is ARF2-modulated, causes eventually enhanced chloroplast development in part by boosting expression of the later, greening phase-associated *GLK1* (Figure 8H) and partly independently of GLKs. This hierarchy of regulation also explains the earlier observation (Waters et al., 2009) that expression of *GLK1* is norflurazon-sensitive, i.e. it occurs only subject to uninterrupted progress through the earlier stages of chloroplast biogenesis. Whether *GLK1* is a direct or an indirect target of GRF5, or indeed of other, functionally redundant products of the *GRF* gene family (Horiguchi et al., 2005; Vercruyssen et al., 2015), may be a subject for future investigation, but the genetic hierarchy remains true regardless.

Chloroplast and leaf development depend on light signalling as a checkpoint, largely through phytochrome-mediated induction of transcriptional drivers (Tepperman et al., 2006; Martín et al., 2016). In *hy1* mutants, defective phytochrome chromophore biosynthesis impairs such light-driven regulation and causes severe leaf development and chloroplast biogenesis defects (Muramoto et al., 1999; Vinti et al., 2000), which we here observe *arf2^sgl2^* dramatically overcomes. Only *GLK2* is substantially light-induced (Tepperman et al., 2006), yet a single *glk2* mutant has no distinct phenotype (Fitter et al., 2002). This means that the rescue of progress through chloroplast biogenesis in *hy1* by *arf2^sgl2^*is a consequence of its impact on the ARF2-GRF5 module not only on *GLK2*, but also on other, GLK-independent drivers of greening. Moreover, the dramatic rescue of the leaf development defects in *hy1* by *arf2^sgl2^* also emphasizes the need for undisturbed chloroplast biogenesis for complete leaf cellular development, as part of a broader, developmental re-interpretation of “retrograde signalling” (Tan et al., 2008; Loudya et al., 2024).

The cellular regulatory logic of chloroplast development is also made apparent by the reduced suppression by *arf2^sgl2^* of the double mutant containing the *glk1*-2 null rather than the initial *glk1* knock-down allele. This led to the observation of a promotion of *GLK1* expression by GRF5, which itself reveals a feed-forward mechanism in the hierarchy of chloroplast biogenesis: action of GRF5 resulting in elevated expression of *GLK1* will drive the greening phase, which in itself will promote *GLK1* expression. Through such a feed-forward mechanism the checkpoint transitions is more likely to be robust and unidirectional. Meanwhile the partial suppression of the null alleles implies that the ARF2-GRF5 module can activate chloroplast development in absence of GLK factors, via other recently found (Frangedakis et al., 2024) or currently unknown regulators.

The extent to which GRF5 directly promotes plastid division is unclear. The increased plastid number per cell resulting from GRF5 overexpression could be only a consequence of the increased cell growth, given the well-established, close correlation between cell size and chloroplast cellular content (Pyke, 1997; Kawade et al., 2013; Larkin et al., 2016). However our data did reveal an increase also in plastid density triggered by *arf2^sgl2^*, and this cannot be explained simply by increases in cell size. What nevertheless remains key is the early action of GRF5, necessary to explain its cell number and growth impact, and the fact that it boosts greening and even chloroplast phenotype (with increased thylakoid stacking) through its action on later components, as part of a chloroplast biogenesis regulatory (direct or indirect) hierarchy (Figure 8H).

In a world in which the competing demands of climate stabilisation, biodiversity protection and food security for an expanding population are ever more apparent, the enhancement of photosynthetic capacity of crops to boost yield ceiling is receiving an increased amount of interest (Long et al., 2015; Eckardt et al., 2023; Croce et al., 2024). Silencing of *ARF2* may provide a potential approach to boost the chloroplast compartment of individual cell types, as would be required for some of the engineering biology approaches currently being explored (Ermakova et al., 2020; Eckardt et al., 2023).

## Acknowledgements

PM was a recipient of a Netaji Subhas International Fellowship from the Indian Council of Agricultural Research, 18(01)/2018-EQR/Edn, and of a fees studentship from Royal Holloway University of London. ELJ and JMH were recipients of BBSRC UK Research and Innovation (UKRI) grant BBP0031171. We are indebted to Ms. Isabelle Williams for her assistance developing the genotyping assays and generating the *arf2^sgl2^ glk1*-2 *glk2*-2 line, and to Prof. Laszlo Bogre and particularly to Dr. Naresh Loudya for immensely skilful technical assistance and many helpful discussions. We are also grateful to Prof. J. Langdale (University of Oxford), Prof. G. Horiguchi (Rikkyo University) and Prof. R. Larkin (Wuhan Agricultural University) for the gift of seed stocks.

## Author contributions

**ELJ** conceptualised the work, with involvement from **PM**, supervised the work and obtained funding**. PM** performed the entirety of the research, contributed to research design, carried out data visualisation and obtained funding. **JMH** co-supervised work and obtained funding. **PM** and **ELJ** co-wrote the manuscript. All authors contributed to the final manuscript.

## Data availability

All data are available in the main text and in Supplemental Figures S1-S7 and Tables S1-S8. Gene accession numbers are listed in Supplemental Table S8.

