## Supplemental Figures and Tables for "An ARF2-GRF5 module regulates chloroplast biogenesis through GLK1 and GLK-independent mechanisms as part of a genetic network"

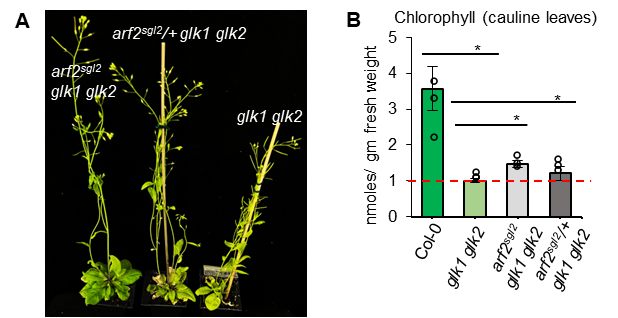

**Supplementary Figure S1, associated to Fig. 2.** *sgl2* is a semi-dominant mutation. (A) Phenotype of F1 progeny along with the parental controls (B) Chlorophyll quantitation of backcross F1 progeny of *sgl2*. The red line indicates chlorophyll concentration in *glk1 glk2*. Analysis was performed using 50-day-old cauline leaves. The error bar represents the means ± SEM (n = 4 replicates). The asterisk symbol represents the significant difference using the Student’s t-test P-value < 0.05 (*).

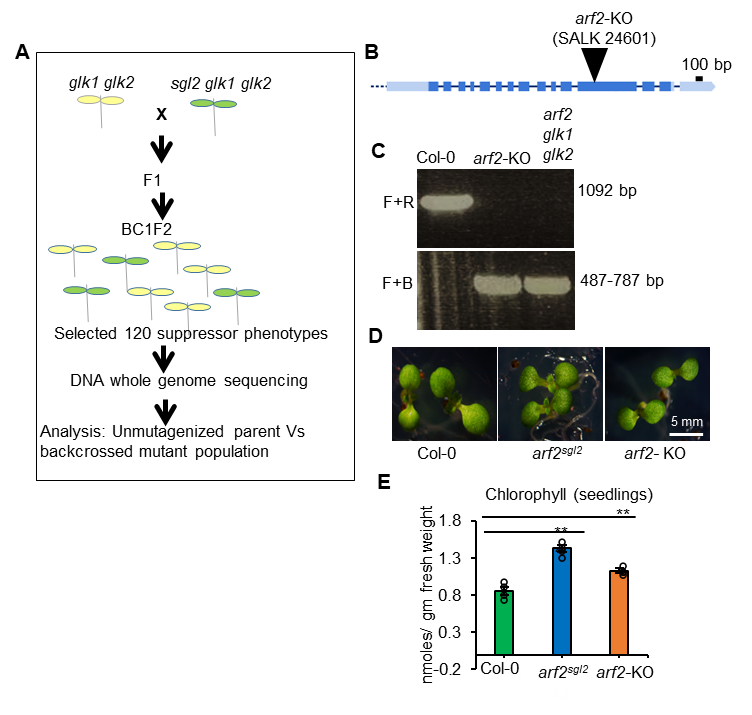

**Supplementary Figure S2, associated to Fig. 3.** Mapping by sequencing strategy. (A) Schematic of generation of a backcrossed population between the suppressor and the parent *glk1 glk2*. In the F2 generation, plants with mutant phenotypes were selected to create the mapping population. The Easymap tool compared the backcross F2 mutant phenotype population with the unmutagenized parent (*glk1 glk2*). (B) The gene structure represents location of T-DNA insertion in *arf2-*KO allele (SALK 24601). (C) Genotyping of *arf2-*KO T-DNA insertional mutant. (D) Phenotype of 7-day old Isolated *arf2^sgl2^* single mutant along with an independent allele, *arf2-*KO. (E) Chlorophyll quantitation in the genotypes mentioned.

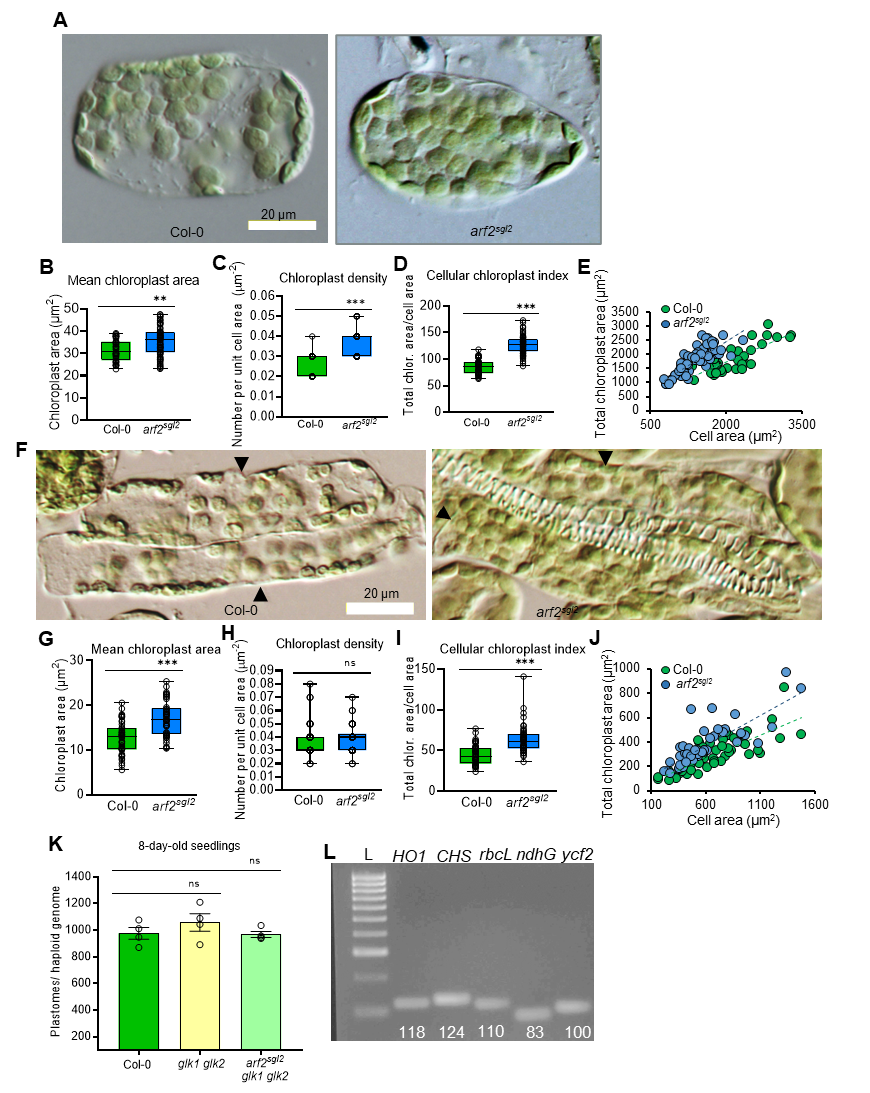

**Supplementary Figure S3, associated to Fig. 4.** Analysis of the cellular chloroplast compartment in mesophyll and bundle sheath cell of isolated *arf2^sgl2^* single mutant. (A) Mesophyll cell of single mutants and WT. (B-E) Individual chloroplast area (B), density (number per unit cell area, C), and total chloroplast area per unit cell area (D, E) in 30-day-old cauline leaves. (F) Images of bundle sheath cells from mutants and WT. Arrowheads indicate individual bundle sheath cells. (G-J) Chloroplast area, chloroplast density and cellular chloroplast compartment (%). Error bars represent ± SEM (n = 3 independent leaves, with a combined total of 40 cells per genotype). Significant differences between mutants and WT are indicated by Student’s t-test P-values ≤ 0.01(*) and ≤0.001 (***). (K and L) Quantitation of number of copies of the chloroplast genome per haploid nuclear genome and gel image showing the PCR amplicons of nuclear and chloroplast targets.

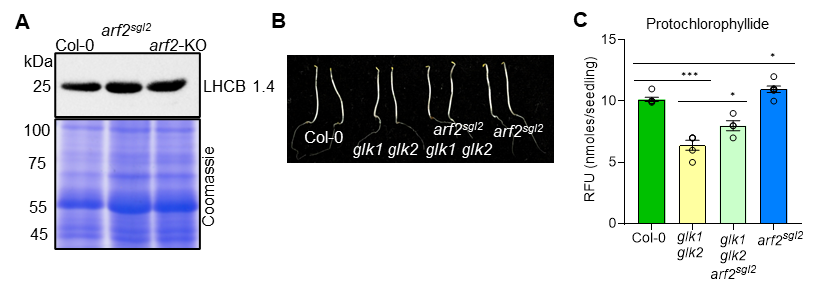

**Supplemental Figure S4, associated to Fig. 5.** Additional phenotypes of *arf2^sgl2^*. (A) 20 µg of total protein of the *arf2^sgl2^* and *arf2-KO* along with WT was separated on a 10% polyacrylamide gel with a PageRuler™ Plus Prestained Protein Ladder. The protein membrane was incubated for 1 hour with anti-LHCB primary and chemiluminescence enzyme-labelled secondary antibody. Protein detection was done using the Chemidoc (Bio-Rad) imaging system. Equal protein loading, of proteins other than chloroplast-contained Rubisco Large subunit (55 kDa), is confirmed in the Coomassie-stained gel. Assay was performed in 30 day old rosette leaves. (B) Morphology of 5-day-old dark grown seedlings. (C) Quantitation of protochlorophyllide content of the mentioned genotypes.

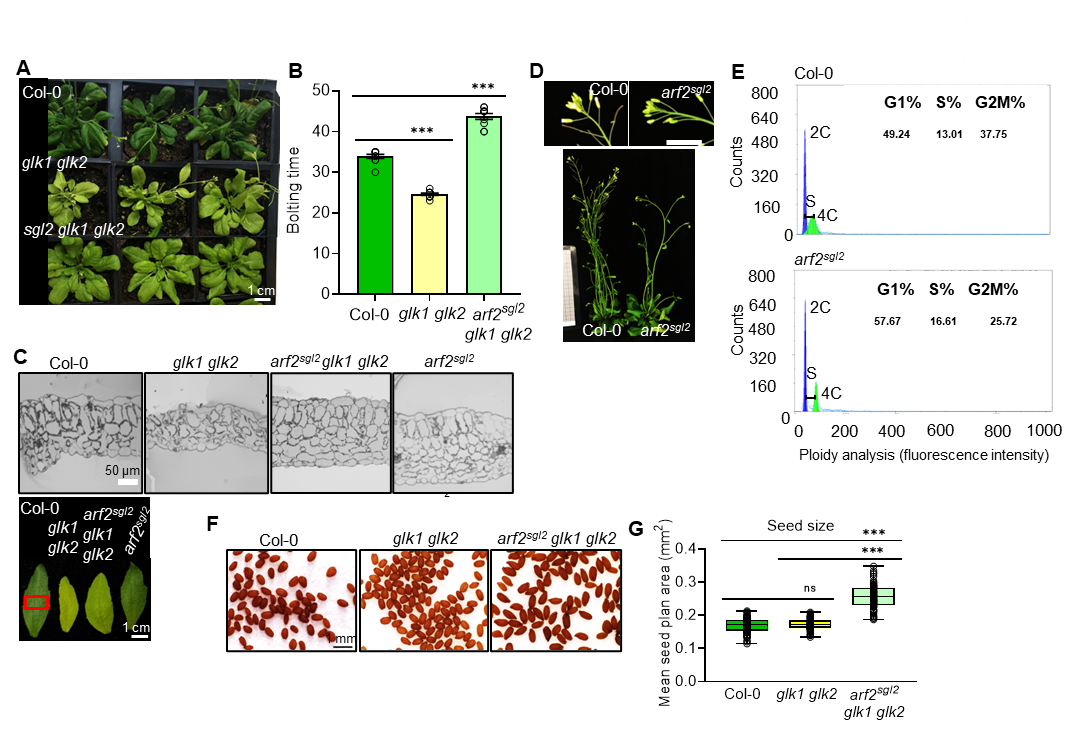

**Supplementary Figure S5, associated to Fig. 6.** Leaf anatomy, ploidy, flowering time and seed size phenotypes of *arf2^sgl2^*. (A and B) Late flowering phenotype, quantified as bolting time. (C) Analysis of leaf anatomy of *arf2^sgl2^.* Light microscopy images showing leaf cellular anatomy in thick transverse sections at 20x magnification. Assay was performed in 30-day-old young rosette leaf sections as indicated by the red box. (D and E) Ploidy analysis in *arf2^sgl2^*. (D) Inflorescence and rosette phenotype. (E) Cell cycle parameters of *arf2^sgl2^* and control. Analysis performed in young buds using 3 biological replicates of the indicated genotypes. Scale bar:1 cm. (F and G) Seed Phenotype of *arf2^sgl2^* and analysis of seed size (plan area). Quantitation was conducted on 60 seeds per genotype. Student’s t-test P-value ≤0.001 (***). Seeds images were captured using a Leica stereomicroscope. Error bars represent ± SEM between replicates (n = 3 independent seed samples).

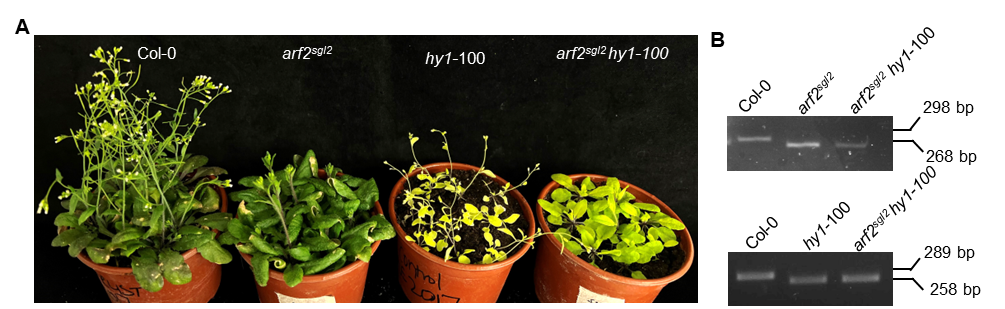

**Supplemental Figure S6, associated to Fig. 7.** *arf2^sgl2^* substantially suppresses the phytochrome-deficient *hy1*-100 mutant. (A) Phenotype of 30-day old *arf2^sgl2^* *hy1-*100 double mutant along with single mutant controls. (B) Genotyping of *hy1*-100 and *arf2^sgl2^* mutations confirming the identity of the double mutant.

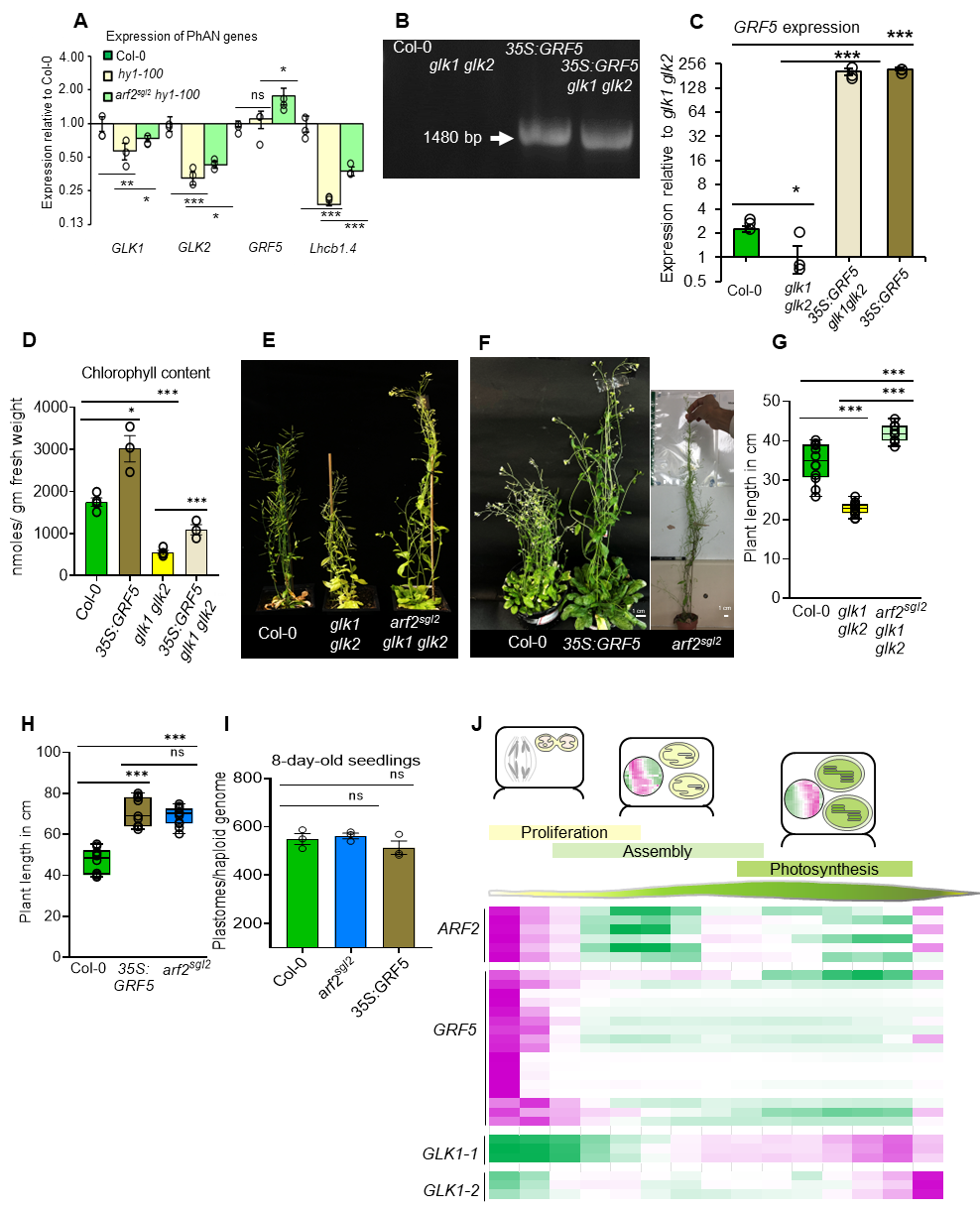

**Supplemental Figure S7, associated to Fig. 8.** Generation and analysis of the *35S:GRF5 glk1 glk2* and *arf2^sgl2^ hy1-100*. (A) Gene expression analysis of *arf2^sgl2^ hy1-100* in in 30-day-old rosette leaves*.* (B) Genotyping of *35S:GRF5* using a forward primer of the 35S promoter sequence from the vector (pB2GW7) and the reverse primer from the *GRF5* gene. (C) Transcript levels of *GRF5* in 7-day-old seedlings. (D) Chlorophyll content of *35S:GRF5 glk1 glk2* and controls, comparable cauline leaves. (E-F) Phenotype of *arf2^sgl2^ glk1 glk2* (E)*, arf2^sgl2^* and *35S:GRF5* (F) and height measurement (G, H) along with their respective controls grown simultaneously. (I) Quantitation of genome copy number, copies of cpDNA per haploid nuclear genome, of *35S:GRF5* and *arf2^sgl2^.* (J) Schematic showing different stages of chloroplast development. Transcript levels of the multiple closest *Triticum aestivum* homologs of *ARF2*, *GRF5,* *GLK1* and *GLK2* in the developing wheat leaf (Loudya et al., 2021). The heatmap represents Z scores. Gene IDs are given in Supplemental Table S8.

**Supplemental Table S1.**  Easymap analysis of highly ranked SNPs list of *sgl2* induced by EMS mutagenesis

| **Position** | **Quality** | **Alt_allele_freq** | **Alt_base** | **aa_pos** | **aa_alt** | **Gene_model** |
| --- | --- | --- | --- | --- | --- | --- |
| 24913807 | 225 | 1 | C-T | 628 | Q-STOP | AT5G62000 |
| 15911945 | 222 | 0.58 | C-T | 135 | P-S | AT5G39760 |
| 6566764 | 222 | 0.57 | C-T | 289 | C-Y | AT4G10630 |
| 7072687 | 222 | 0.55 | C-T | 86 | P-L | AT4G11740 |
| 2988905 | 222 | 0.55 | G-A | 440 | R-K | AT2G07190 |
| 7298328 | 222 | 0.55 | C-T | 174 | S-F | AT4G12270 |
| 10455564 | 222 | 0.54 | G-A | 254 | S-N | AT1G29860 |
| 9292182 | 222 | 0.51 | C-T | 283 | T-M | AT4G16480 |
| 1362087 | 222 | 0.49 | G-A | 352 | E-K | AT5G04720 |
| 13334181 | 222 | 0.49 | C-T | 19 | G-E | AT4G26370 |

The highlighted gene ID is the most likely candidate

**Supplemental Table S2.** List of genotyping and cloning primers

| **Target** | **Forward/ reverse** | **Sequence** |
| --- | --- | --- |
| *glk1* | Border | GATCCGACACTCTTTAATTAACTGACA |
|  | Forward | GGTATTTTGGGTTCGGGTTT |
|  | Reverse | GTTGTTACTGATCCGATTGTTCTTG |
| *glk2* | Border | GTTTTGGCCGACACTCCTTA |
|  | Forward | CATGTCAGTATCCACCAACACA |
|  | Reverse | TCCGATGTGACCTATATTTC |
| *arf2*-KO | Border | ATTTTGCCGATTTCGGAAC |
|  | Forward | ATGAAGATTTTGCGAACCATG |
|  | Reverse | TTACACAGATTTGCTCTCCGG |
| *ARF2*_gDNA | Forward | GGGGACAAGTTTGTACAAAAAAGCAGGCTTCGATGCGGCAGAGATGAAA |
|  | Reverse | GGGGACCACTTTGTACAAGAAAGCTGGGTCGTAGCTAGAAACATCCGAA |
| *grf5* SALK line | Forward | CACCAATCCCGTAAGCCCTA |
|  | Reverse | GCGCCCAAAACAAAACACTG |
| *glk1*-2 (CRISPR) | Forward | AAGAGATGGTTGCGACGGAG |
|  | Reverse | ACGACTTCTTCACCTTTCCCC |
| *glk2*-2 (CRISPR) | Forward | AACTGTTTCTCCGGCTCCAG |
|  | Reverse | ATTGCTCCACCGCTTGTACA |

**Supplemental Table S3.** dCAPS primers and restriction enzyme used to genotype *sgl2* and *hy1*-100 point mutations

| **Genotype** | **Primer** | **Forward/ reverse** | **Sequence** | **Enzyme digestion** | **Band size (bp)** |
| --- | --- | --- | --- | --- | --- |
| *sgl2* | dCAPS_*sgl2* | Forward | TATTATGAGGAAGTGGTCAATGCTCAA GCT | HindIII | Col-0 (298)  *sgl2* (268) |
|  |  | Reverse | TCTGGAATGGTCTTCCCTGTT |  |  |
| *hy1*-100 | dCAPS_*hy1-100* | Forward | TCTTGGATTGAGTTGTTGGTTGTGGTGT | ApoI | Col-0 (289)  *hy1-100* (258) |
|  |  | Reverse | TTTCCAGCCCCGTGTTCTTGAACTCGGAAT |  |  |

**Supplemental Table S4.** List of antibodies used and their dilutions

| **Antibody** | **Dilution** |
| --- | --- |
| LHCB1 AS01 004 | 1:2000 |
| PsbO AS06 142-33 | 1:2000 |
| IRDye secondary antibody | 1:5000 |
| HRP-conjugated secondary | 1:15000 |

**Supplemental Table S5.** List of cloning vectors and primers

| **Genotyping primers after BP reaction** | | | |
| --- | --- | --- | --- |
| pDONR201  (ENTRY) | Forward | TCGCGTTAACGCTAGCATGGATCTC | |
| ARF2clo_gDNA_R | Reverse | GTAGCTAGAAACATCCGAA | |
| **Genotyping primers for plant transformants** | | | |
| PKGW1  (DESTINATION) | Neo-F | Forward | AATATCACGGGTAGCCAACG |
| ARF2_gDNA_R1 | gDNA | Reverse | TAGTCCACTCATACAGCTGGC |

**Supplemental Table S6.** Paraments used for thylakoid development

| **Parameters** | **Calculation** |
| --- | --- |
| Chloroplast area (µm^2^) | - |
| No. of grana stacks | Grana stacks counted only having 3 ≥ lamellae |
| Mean width of 10 grana stacks (µm) | - |
| Thylakoid distribution | No. of grana stacks (3 ≥ lamellae) X Mean width of 10 grana stacks (µm) |
| Thylakoid density (µm^-1^) | Thylakoid distribution/chloroplast area |
| Analysis was done with ≥ 20 chloroplasts | |

**Supplemental Table S7.** List of RT-qPCR primers

| Target | Direction | Sequence |
| --- | --- | --- |
| GLK1 | Forward | TTGGGTCTCCGATTCTCCCTAT |
|  | Reverse | GCAACTGGCGGTGCTCTAAAT |
| GLK2 | Forward | ACCGTACTGGCATCAGCAAC |
|  | Reverse | TGAATGTCGATGGGAGGATT |
| UBQ10 | Forward | GGAGGATGGTCGTACTTTGG |
|  | Reverse | TCCACTTCAAGGGTGATGGT |
| LHCB1.2 | Forward | CCGATCCAGTCAACAACAAC |
|  | Reverse | TCAAACCATCACATACAACCTTC |
| GRF5 ([Vercruyssen](https://pubmed.ncbi.nlm.nih.gov/?term=Vercruyssen%20L%5BAuthor%5D) et al., 2015) | Forward | TCAGTTCAATGTCTTAGCCTCTGC |
|  | Reverse | CCCAACTCCTCCAACTCTCTCC |
| HEMA1 | Forward | GCTTCCGCAGTCTTCAAACG |
|  | Reverse | CCAGCGCCAATTACACACATC |
| CAO | Forward | TCGAAAGCACACCAACAGGTTC |
|  | Reverse | GGGCTAACTTGTCTAAAGCAGTCG |
| GUN4 | Forward | CTGCCGTTTCAACCACAAACGC |
|  | Reverse | ACGTCGAATATGGTCGCGGTTTC |
| CHLH | Forward | TGGTAGAGAGACAGAAGCTCG |
|  | Reverse | CCAAAGAACCTGCCCAAGAG |

**Supplemental Table S8.** Identities of the *ARF2* and *GRF5* genes in the Arabidopsis genome and of their homologues present in the *Triticum aestivum* genome, whose expression is displayed in Supplemental Figure S7.

| **Gene** | **Closest homolog in Arabidopsis** |
| --- | --- |
| **ARF2** |  |
| TraesCS3A02G442000 | AT5G62000 |
| TraesCS3A02G449300 | AT5G62000 |
| TraesCS3B02G475800 | AT5G62000 |
| TraesCS3B02G486000 | AT5G62000 |
| TraesCS3D02G434700 | AT5G62000 |
| TraesCS3D02G442000 | AT5G62000 |
| **GRF5** |  |
| TraesCS2A02G238700 | AT3G13960 |
| TraesCS2A02G435100 | AT3G13960 |
| TraesCS4A02G434900 | AT3G13960 |
| TraesCS6A02G269600 | AT3G13960 |
| TraesCS6A02G335900 | AT3G13960 |
| TraesCS6B02G296900 | AT3G13960 |
| TraesCS6B02G366700 | AT3G13960 |
| TraesCS6D02G245300 | AT3G13960 |
| TraesCS6D02G315700 | AT3G13960 |
| TraesCS7A02G049100 | AT3G13960 |
| TraesCS7A02G165600 | AT3G13960 |
| TraesCS7B02G070200 | AT3G13960 |
| TraesCS7D02G044200 | AT3G13960 |
| TraesCS7D02G166400 | AT3G13960 |
| TraesCS2B02G458400 | AT3G13960 |
| TraesCS2D02G246600 | AT3G13960 |
| TraesCS2D02G435200 | AT3G13960 |
| **GLK1_1** |  |
| TraesCS7A02G339800 | AT2G20570 |
| TraesCS7B02G251400 | AT2G20570 |
| TraesCS7D02G347500 | AT2G20570 |
| **GLK1_2** |  |
| TraesCS3A02G161000 | AT2G20570 |
| TraesCS3B02G191600 | AT2G20570 |
| TraesCS3D02G168200 | AT2G20570 |
